# The impact of α-synuclein aggregates on blood-brain barrier integrity in the presence of neurovascular unit cells

**DOI:** 10.1101/2022.08.18.504449

**Authors:** Hamdam Hourfar, Farhang Aliakbari, Shabboo Rahimi Aqdam, Zahra Nayeri, Hassan Bardania, Daniel E. Otzen, Dina Morshedi

## Abstract

The role of the blood-brain barrier (BBB) is to control trafficking of biomolecules and protect the brain. This function can be compromised by pathological conditions. Parkinson’s disease (PD) is characterized by the accumulation of α-synuclein aggregates (αSN-AGs) such as oligomers and fibrils, which contribute to disease progression and severity. Here we study how αSN-AGs affect the BBB in *in vitro* co-culturing models consisting of human brain endothelial hCMEC/D3 cells alone and co-cultured with astrocytes and neurons/glial cells. When cultivated on their own, hCMEC/D3 cells were compromised by αSN-AGs, which decreased cellular viability, mitochondrial membrane potential, wound healing activity, TEER and permeability parameters, as well as increased the levels of ROS and NO. Co-culturing of these cells with activated microglia also increased BBB impairment according to TEER and systemic immune cell transmigration assays. In contrast, hCMEC/D3 cells co-cultured with astrocytes or dopaminergic neurons or simultaneously treated with their conditioned media showed increased resistance against αSN-AGs. Our work demonstrates the complex relationship between members of the neurovascular unit (NVU) (perivascular astrocytes, neurons, microglia, and endothelial cells), αSN-AGs and BBB.

**Graphical Abstract:** 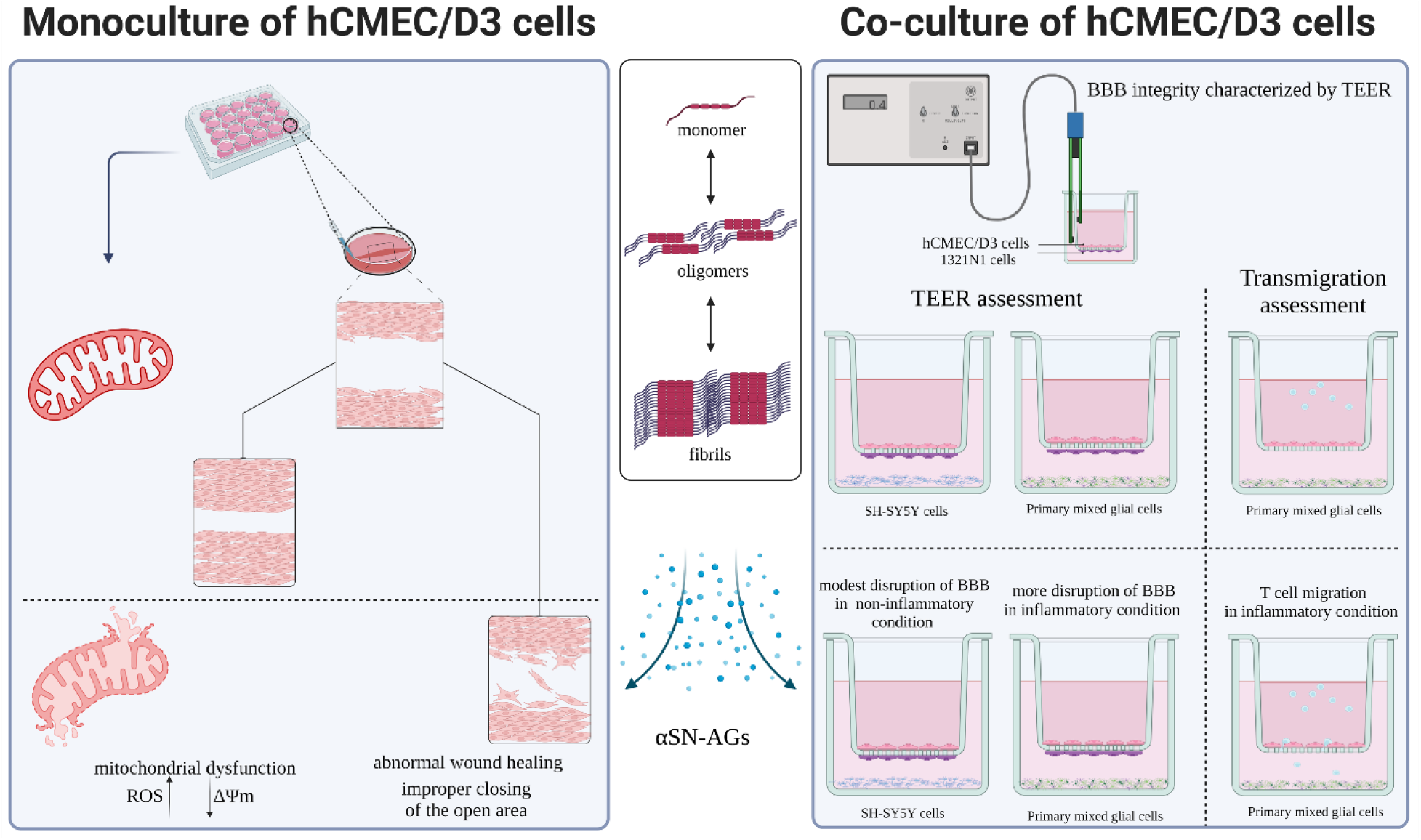

## Background

The human brain, the most intricate organ in the body, is protected and maintained by a highly sophisticated system known as the blood-brain barrier (BBB) [1]. Although different studies have shown that BBB integrity can be altered or disrupted by various pathological conditions, there is still debate about the BBB’s role in the onset or progression of neurodegenerative diseases (NDs) such as Parkinson’s disease (PD) [2–4]. NDs may compromise BBB integrity through reactive oxygen species (ROS), nitric oxide (NO), matrix metalloproteinases (MMPs), inflammatory cytokines, and immune cell extravasation. These can lead to breakdown of the BBB, mainly by downregulation of tight junction proteins and perturbation of the TEER (trans-endothelial electrical resistance) and permeability parameters, which consequently promote neuro-inflammation and also neurodegeneration [5]. In support of this, PD patients have been shown to undergo significant BBB abnormalities, such as disruption of its integrity and decreased P-glycoprotein function [6, 7]. However, the exact role of BBB in PD is unclear. Some clues may be found by considering the etiology of PD, in particular the formation of intracellular inclusions known as Lewy bodies and Lewy neurites, which are rich in amyloid aggregates formed by the protein α-synuclein (αSN-AGs) [8, 9]. Other toxic species include soluble oligomeric versions of αSN which are even more cytotoxic than amyloid fibrils on a per-mass basis [10]. In contrast to the intense focus on the relationship between PD and αSN aggregation, there are few studies on the impact of αSN and αSN-AGs on BBB integrity and functionality. The BBB consists of specified endothelial cells which interact with astrocytes, pericytes, neurons, and microglial cells, collectively known as the neurovascular unit (NVU) [1, 11]. The role of these cells on BBB integrity and its function in pathological conditions is important to consider.

αSN-AGs may induce inflammatory responses in the brain that may subsequently affect BBB integrity. αSN directly interacts with Fcγ and toll-like receptors (TLRs) and induces nuclear factor-kappa B (NF-kB) and mitogen-activated protein kinase (MAPK) signal pathways or indirectly induces expression of the matrix metalloproteinases (MMPs) [12]. Subsequently, microglial inflammatory signals and inflammatory-related cytokines can be induced [12, 13]. Inflammation is common in PD [14], and it is likely to compromise BBB integrity in different ways. Disruptive changes include endothelial damage such as apoptosis, membrane abnormalities, and mitochondrial dysfunction; impairment of tight junction integrity; degradation of the substance that coats the luminal surface of blood vessels (glycocalyx), and astrocyte destruction. Non-disruptive changes include induction of cytokines, prostaglandin E_2_ production, dysregulation of transporters, and cellular transmigration [15]. Besides these indirect effects of αSN-AGs on the BBB integrity through inflammation, it is also important to evaluate how αSN-AGs can directly affect BBB since this may be an important aspect of their neurotoxicity.

An important aspect in this regard is the primary origin of αSN aggregates. αSN is expressed in various tissues, in particular the brain and hematopoietic system cells [16–18]. It is possible that early toxic aggregated species of αSN are produced in peripheral organs and cross BBB. αSN is known to cross the BBB in both directions [19, 20], although recent work shows that monomeric αSN, taken up in a clathrin-dependent pathway, is transported in a polarized fashion, largely in a luminal-abluminal direction [21]. On the other hand, oligomeric αSN is transported across the BBB to a much smaller extent [21], and transport of fibrillar species is likely to be even less efficient. Thus, αSN aggregates could also be formed as a secondary response to neuronal deterioration and αSN clearance. If the BBB cannot dispose properly of protein aggregates, the inflammatory response induced by brain’s glial cells could play a considerable role in the progression of neurodegenerative disease [22–24], as reported for Alzheimer’s disease [25].

Given the limited insight into the effects of αSN-AGs on BBB, we firstly investigated the influence of different αSN-AGs (formed after different periods of incubation) on brain endothelial cell viability, ROS and NO content, as well as mitochondrial membrane potential (ΔΨm). Furthermore, BBB parameters such as TEER and permeability values, as well as wound healing properties were examined. The study was expanded by co-culturing brain endothelial cells with astrocytes, dopaminergic neurons, and activated glial cells. We also explored the transmigration of systemic immune cells. Finally, we compared the interactions of αSN aggregates with brain endothelial cells, non-brain endothelial cells, and astrocytes.

## Materials and methods

### Materials

2ʹ,7ʹ- dichlorodihydrofluorescin diacetate (DCFH-DA), 3-(4,5-dimethylthiazol-2-yl)-2,5-diphenyltetrazolium bromide (MTT), collagen, fluorescein isothiocyanate–dextran (FITC-Dextran 70), FITC, Rhodamine 123, 4′,6-Diamidino-2-phenylindole dihydrochloride (DAPI), Griess reagents, Thioflavin T (ThT) and Congo red (CR) were all from Sigma-Aldrich. SH-SY5Y cells overexpressing human αSN were generously provided by Kostas Vekrellis and Stefanis Leonidas from the Biomedical Research Foundation of the Academy of Athens (BRFAA), Greece [26]. Roswell Park Memorial Institute (RPMI) 1640 medium, DMEM/F-12 (Dulbecco’s Modified Eagle Medium/Nutrient Mixture F-12) and DMEM high glucose were from GibcoBRL (Life Technologies, Paisley, Scotland). Fetal bovine serum (FBS) was from Biosera (Tehran, Iran). Transwell 24-well tissue culture inserts (0.4 μm pore size, polyester membrane) and (8.0 μm pore size, polycarbonate membrane) were from Corning Costar (Corning Inc., Coring, NY, USA).

### Methods

All fluorescence measurements were carried out using a Cary Eclipse fluorescence spectrophotometer (Agilent, Santa Clara, CA).

### Induction and assessment of αSN-AGs formation

#### Preparation of αSN-AGs

Human αSN protein was expressed in *Escherichia coli*, strain BL21(DE3) (Novagen, Madison, Wis., USA), and purification of protein was carried out as described previously [27]. 70 μM αSN was dissolved in the fibrillation solution (phosphate buffered saline (PBS) containing 0.2 mM phenylmethylsulfonyl fluoride, 1 mM ethylenediaminetetraacetic acid and 0.05 mM NaN_3_. Before incubation, αSN solution was centrifuged at 16000×g for 10 min and then incubated at 37 °C, with 300 rpm shaking in 96-well plates containing glass beads [28].

#### ThT fluorescence assay

Thioflavin T (ThT) is a fluorescent dye which gives a strong fluorescence signal when bound to amyloid fibrils [29]. To monitor fibrillation, 10 μL of the incubated αSN solution was collected at different times during fibrillation and added to 490 μL of ThT solution (30 μM of ThT dissolved in Tris-buffer (10 mM), pH: 8). Intensity of ThT fluorescence was measured with excitation at 440 nm and emission spectra in the range of 450 to 550 nm using excitation and emission slit widths of 5 and 10 nm, respectively.

#### Congo red (CR) absorbance assay

Congo red (CR) undergoes an absorption change in the visible region when bound to amyloid fibrils *in vitro* [30]. To this end, 10 μL of the incubated αSN solution at different times during fibrillation was added to 490 μL of the CR solution (80 μM in a buffer consisting of 5 mM KH_3_PO_4_, 150 mM NaCl, pH 7.4) and incubated in the dark for 5-10 min. Absorption spectra (400–600 nm) were recorded using a PGT80+UV– Visible spectrometer (Leicestershire, England).

#### Atomic force microscopy

Atomic force microscopy (AFM) imaging is a single molecule technique that has been widely employed to visualize the nano-mechanical properties and structure of amyloid fibrils [31]. To probe the structural details of αSN-AGs during the fibrillation process, 3-, 12-, and 24-h αSN-AGs were collected. The samples were diluted 20 times with the filtered deionized water, and 10 μL of each sample was deposited on a freshly cleaved mica sheet and dried at room temperature. All images were taken using a Veeco scanner equipped with a silicon probe (CP), and imaging was done in noncontact mode.

#### Fluorescent staining of αSN-AGs

To monitor different stages of αSN fibrillation by fluorescence microscopy [32], 15 μL of the incubated αSN (3-, 12-, and 24-h αSN-AGs) was added to the 15 μL of 500 μM ThT and the mixture was incubated in the dark for 5 min at room temperature. Thereafter, samples were spread onto microscopy slides and fluorescence microscopy images were recorded using a Nikon Ti Eclipse inverted microscope (Nikon, Tokyo, Japan).

#### Cell culture

The human brain endothelial cell line (hCMEC/D3) is a well-known endothelial cell line employed for different *in vitro* BBB models [33–35]. The cells were cultured in RPMI-1640 medium supplemented with 5% FBS and incubated at 37 °C with 95% humidity, 5% CO_2_.

Wild type SH-SY5Y (WT SH-SY5Y) and Tet-Off SH-SY5Y cells which expressed αSN in the absence of doxycycline (O-αSN SH-SY5Y) [26], 1321N1 cells (a human astrocytoma cell line) as well as Jurkat cells (an immortalized line of human T lymphocyte cells), HUVEC cells (umbilical vein endothelial cell) and primary mixed glial cells were cultured in their specific media including DMEM-F12 (WT SH-SY5Y and O-αSN SH-SY5Y cells), RPMI-1640 (1321N1 and Jurkat cells) and high glucose DMEM (primary mixed glial cells), with 10% fetal bovine serum (FBS) and incubated at 95% humidity, 5% CO_2_ and 37 °C. O-αSN SH-SY5Y cells were cultured with (2 μg/ml) or without doxycycline supplementation to suppress or induce αSN expression, respectively [26]. Mixed primary glial cells were obtained from the brains of Wistar rat pups (no older than 72 h) and cultured as described [36]. To obtain primary microglia-rich mixed glial cultures, media were replaced with fresh media after 2-3 days, and cells were trypsinized and seeded on 24-well plates 14-21 days after their initial seeding. All procedures for animal management, euthanasia and surgery were complied with the guidelines of the National Institute of Genetic Engineering and Biotechnology Ethics Committee (ethics number: IR. NIGEB. EC.1398.10.18. A).

#### Assays to measure the effects of αSN-AGs on hCMEC/D3 cells and BBB

The effects of αSN-AGs on hCMEC/D3 cells and BBB were measured through cell metabolic activity (MTT assay), intracellular ROS measurement, NO analysis, mitochondrial membrane potential (ΔΨm) assay, cell migration and cell-cell and cell-ECM interactions (scratch assay), measurement of TEER value and the rate of permeability (using FITC-Dextran), measurement of transmigration of Jurkat cells across the BBB and the interaction of various αSN-AGs with the cells. Cells were cultured as mentioned above, and 96-well plates were used for assessment of MTT, NO analysis, and intracellular ROS, and 24-well plates for mitochondrial membrane potential (ΔΨm) assay, measurement of the interaction of various αSN-AGs with the cells, the inflammation response of the primary mixed glial cells, and the scratch test. TEER and permeability measurements and transmigration of Jurkat cells across the BBB were performed on transwell 24-well tissue culture inserts 0.4 μm pore size and 8.0 μm pore size, respectively.

#### MTT assay

This assay was designed to measure the cell viability. Living cells have active mitochondria with dehydrogenases that convert tetrazolium MTT dye to insoluble purple formazan crystals. Cell suspensions were seeded on 96-multiwell plates at a density of 2×10^4^ cells/cm^2^ in 200 μL medium. Then the cells were treated with 10% v/v of fresh αSN and different αSN-AGs pre-incubated for 3, 7, 12, and 24 h under fibrillation conditions, as well as with different concentrations of 12- and 24-h αSN-AGs (1, 5, 10, and 20 % v/v). The media were then replaced after 24 h with fresh media containing MTT (10% (v/v) of stock solution, 5 mg/mL), and incubated for 4 h. Then, 100 μL DMSO was added to dissolve formazan crystals, and the absorbance was measured at 570 nm (microplate spectrophotometer, Epoch 2, BioTek company, Gen5 software, USA). This process was repeated for 1321N1 and SH-SY5Y cells treated with 12-h αSN-AGs (10% v/v). Conditioned media (-Cond-Med) derived from 1321N1 and SH-SY5Y cells in the absence or presence of 12-h αSN-AGs (5% v/v) were added to hCMEC/D3 cells in the presence of 12-h αSN-AGs (10% v/v).

#### Measurement of intracellular ROS

Dichlorodihydrofluorescein diacetate (DCFH-DA) is a cell permeable compound which converts to its fluorophore derivative, 2’7’-Dichlorodihydrofluorescein (DCF) in the presence of ROS. To measure this, cells were seeded on 96-multiwell plate at a density of 1.5×10^4^ cells/cm^2^ and, after 24 h, treated with 12-h αSN-AGs or 24-h αSN-AGs (10% v/v) and incubated for another 6 h. Then, the media were replaced with a fresh one without FBS, containing 15 μM DCFH-DA, and incubated in the dark for 45 min. To measure the intensity of DCF, cells were trypsinized and DCF fluorescence intensity was measured with excitation at 495 nm and emission at 500-550 nm, with an emission peak at 520 nm and a slit width of 5 nm for both excitation and emission.

#### Assessment of the mitochondrial membrane potential

The potential of the mitochondrial membrane (ΔΨm) was determined using rhodamine-123 (RH-123), which is taken up by active mitochondria with a negative membrane potential. RH-123, like other fluorophore dyes, has the ability to self-quench in high concentrations (1-10 μM). Therefore, a high loading concentration of RH-123 leads to a critical concentration for self-quenching which will decrease fluorescence. A decrease in ΔΨm due to depolarization will release RH-123 from mitochondria to the cytoplasm, leading to a rapid increase in fluorescence intensity. However, dysfunctional mitochondria will lead to a lower ΔΨm, a lower accumulation of RH-123 and a reduction in the post-quenching, fluorescence intensity increase, while the continued loss of ΔΨm leads to a decrease in fluorescence intensity [37, 38]. 10^5^ cells/cm^2^ were cultured on 24-well plates and, after reaching 90% confluence, RH-123 was added at a final concentration of 10 μM to all treatment groups, and after 45 min, the plates were washed twice with PBS, trypsinized, and the cells were resuspended in PBS. The fluorescence emission spectra were measured in the range of 495 to 570 nm (with a peak at 525.07) with the excitation at 485 nm with a slit width of 10 nm for both excitation and emission. Treatments were done in 6 groups (summarized in **Table 1**): 1 group without any treatment (control); 4 groups containing 12-h αSN-AGs (10% v/v) with 0, 0.5, 3, and 6 h duration of treatment before trypsinization in which 20 μM verapamil was added to each group 15 min before adding RH-123 to partially inhibit the activity of the ATP-dependent efflux pump P-gp and thus accelerate RH-123 uptake into the cells; and finally one group containing 50 μM verapamil for complete inhibition of P-gp as a healthy control with 1 h incubation time.

**Table 1.** Overview of treatment groups for assessment of the mitochondrial membrane potential

| Groups | Treatment | Duration of treatment (min)<br>before trypsinization |
| --- | --- | --- |
| **Group 1** | --- | --- |
| **Group 2** | 12-h αSN-AGs (10% v/v) | 0 |
| **Group 3** | 12-h αSN-AGs (10% v/v) | 30 |
| **Group 4** | 12-h αSN-AGs (10% v/v) | 180 |
| **Group 5** | 12-h αSN-AGs (10% v/v) | 360 |
| **Group 6** | Verapamil (50 μM) | 60 |

12-h αSN-AGs α-synuclein incubated for 12 h under fibrillation condition

#### DAPI staining (Nuclear morphology assay)

The nucleus is stained with DAPI, which is a membrane-permeable dye and stains nucleic acids [39]. hCMEC/D3 cells were seeded on 24-well plates with 5×10^4^ cells/cm^2^ confluence and after 24 h treated with 12-h αSN-AGs or 24-h αSN-AGs (10% v/v) and incubated for another 24 h. Then, the cells were washed with PBS and stained with DAPI (1 μg/mL) for 10 min in the dark. After washing with PBS, fluorescence microscopy images were captured by a Ceti inverso TC100 microscope (Medline Scientific, Oxon, UK).

#### Scratch test to assess wound healing capability of hCMEC/D3 cells

The scratch wound assay is a common method to estimate the ability of the cells to migrate, which is a very important function for healing and repairing wounds. hCMEC/D3 cells were seeded on 24-well plates with 10^5^ cells/cm^2^ confluence. A horizontal straight line was scratched on the cell-coated surface using a sterile pipette tip. After twice washing with PBS, the medium with no FBS was added; a group of cells was treated with 12-h αSN-AGs (5% v/v), a group with 24-h αSN-AGs (5% v/v), and the last group was control without any treatment. Cellular migration was analyzed after 24 h and 48 h of treatment. The area of the scratched field was quantified using ImageJ software and the percentage of scratched field area and scratched field closure were calculated according to the following equations [40, 41]:

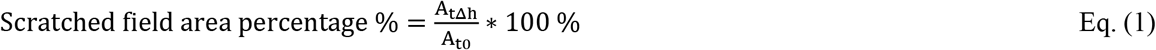

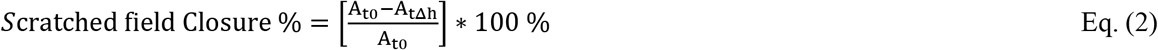

where A_t0_ is the area of the scratch wound at time zero and A_tΔh_ is the area of the scratch wound at time h (24 or 48).

The same procedures were reiterated for HUVEC (umbilical vein endothelial cells as non-special BBB endothelial cells) and 1321N1 cells (astrocytoma cells as non-endothelial cells) to compare their response to that of hCMEC/D3 cells.

### Verify the BBB integrity in different culturing models

#### Monoculture model

The apical side of the inserts was coated with collagen for 24 h, before seeding of hCMEC/D3 cells at a density of 5×10^4^ cells/cm^2^ on the apical side of the inserts for both TEER and trans-endothelial permeability parameters (Fig. 1a). The coated insert without cells was used as a negative control (blank) for TEER and permeability assays. After forming a uniform monolayer of the cells on the insert with 100% confluence and the TEER reaching to a constant value, all treatments described in following sections were applied.

**Fig. 1.**
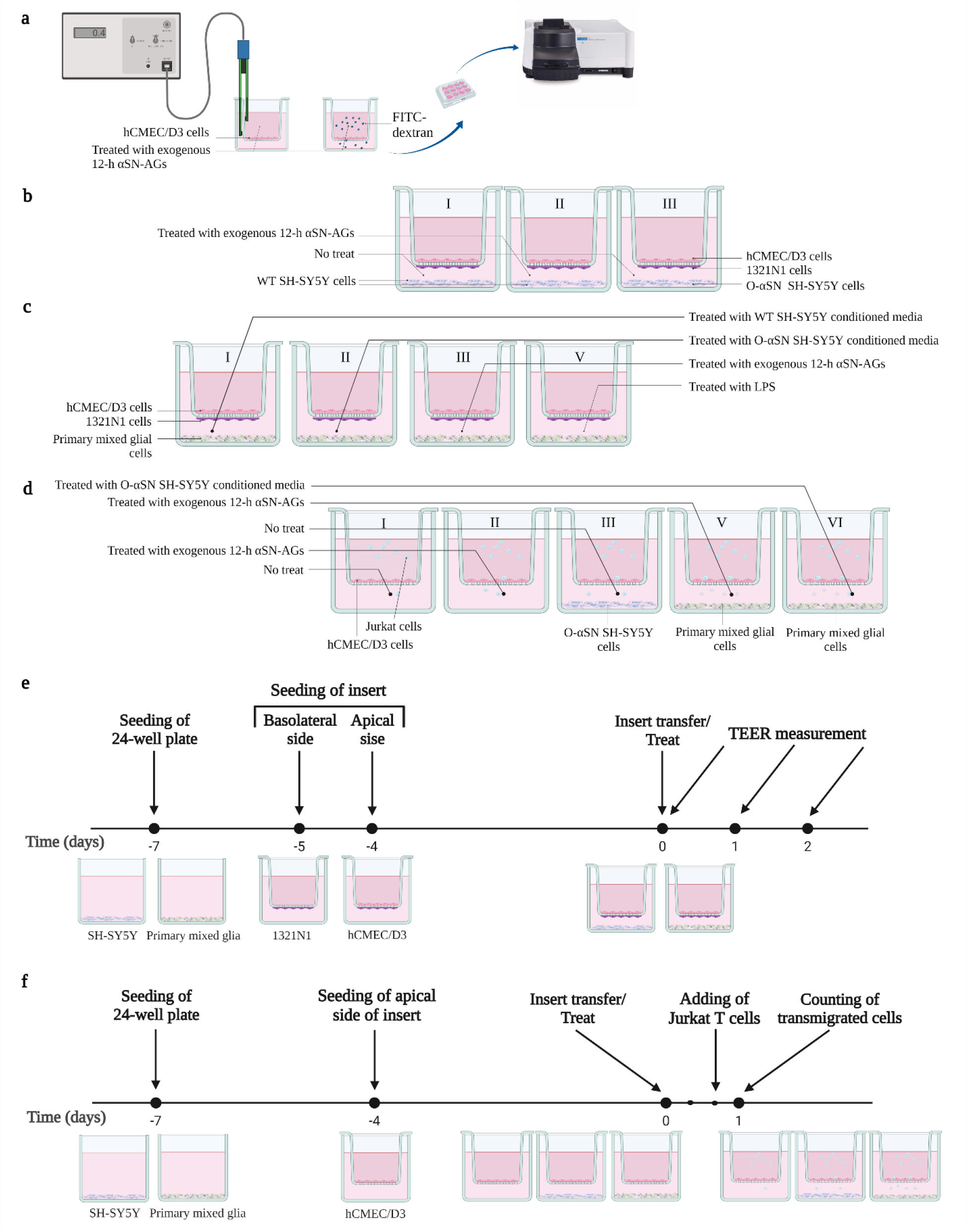
Schematic illustration of the transwell system and experimental designs for co-culture models. (a) hCMEC/D3 cells were cultured as monolayer on the apical side of insert, and TEER and permeability parameters were assessed in the monoculture system in the presence and absence of αSN. (b, c) Transwell showing triple co-culture system containing hCMEC/D3, 1321N1 on the apical and underside of the insert, respectively, and WT SH-SY5Y/O-αSN SH-SY5Y cells or primary mixed glial cells on the bottom of plates. (d) Transwell showing co-culture system containing hCMEC/D3 on the apical side of the insert and no cell or O-αSN SH-SY5Y cells or primary mixed glial cells on the plate bottom. Treatments/cell type of the lower compartment of each transwell are as follows; (bI) no treat/WT SH-SY5Y, (bII) exogenous 12-h αSN-AGs/WT SH-SY5Y, and (bIII) no treat/O-αSN SH-SY5Y; (cI) WT SH-SY5Y conditioned media/Primary mixed glia, (cII) O-αSN SH-SY5Y conditioned media/Primary mixed glia, (cIII) exogenous 12-h αSN-AGs/Primary mixed glia, (cIV) LPS/Primary mixed glia; (dI) no treat/no cell (control group), (dII) exogenous 12-h αSN-AGs/no cell, (dIII) no treat/O-αSN SH-SY5Y, (dIV) O-αSN SH-SY5Y conditioned media/Primary mixed glia, (dV) exogenous 12-h αSN-AGs/Primary mixed glia. Timeline graph of entire experiment for measurement of TEER (e) and transmigration assay (f). The first day of treatment is considered as time zero. (Created with BioRender.com)

#### TEER measurements

Transendothelial electrical resistance (TEER) is a determinant of the activity of the tight junctions [42]. After full coating of transwell inserts with a monolayer of hCMEC/D3, cells were treated with 12-h αSN-AGs (5% v/v) for 16 h and TEER was measured using the EVOM2 apparatus with STX2 chopstick electrodes (World Precision Instruments, USA).

Note: Before and after every measurement, the media of the apical and basolateral compartments were replaced by fresh ones, after twice washing with PBS. The TEER_(tissue)_ or the unit area resistance of the cell monolayer, was calculated by equation (3):

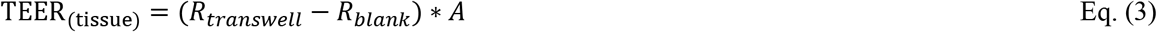

where R_transwell_ is the total resistance (Ω) of the cell monolayer, R_blank_ is the resistance of the coated blank filter (without cell), and A is the surface area of the filter (0.33 cm^2^).

#### Measurement of the permeability parameter of the endothelial cell monolayer

By measuring the paracellular permeability of the specific fluorophore compounds, especially FITC-dextran, the presence of the tight junctions and the strength of the impermeability of the endothelial monolayer can be determined.

After formation of the monolayer, cells were treated with 12-h αSN-AGs (5% v/v) for 16 h; the inserts were transferred to 24 well plates containing fresh media with FBS 1% and washed twice with PBS. The media of the inserts were replaced with fresh media containing FBS 1% and 70kDa FITC-dextran at a final concentration of 100 μg/mL. Then samples were collected from lower compartments at some interval of time (to define the permeability rate from the graph permeability according to previous studies [43]) and fresh pre-warmed media were replaced. The fluorescence intensity was measured with excitation at 495 nm and emission spectra in the range of 505 to 550 nm (peak at 516.86 nm) using excitation and emission slit widths of 5 and 10 nm, respectively. *In vitro* permeability (Pe) was calculated as follows:

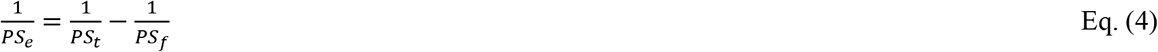

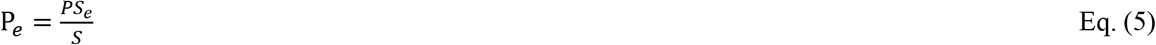

where PS_e_ is the permeability surface area product which is calculated according to the slope of the graph (obtained by average cumulative cleared volume (CL) over time) for cell monolayer on the inserts (total PS, PS_t_), and coated blank filter insert (without cells) (PS_f_). S is the filter surface area (0.33 cm^2^).

### Triple co-culture models

#### Experimental design of co-culturing groups

For triple co-culture model with dopaminergic neurons, WT SH-SY5Y or O-αSN SH-SY5Y cells (5×10^4^ cells/cm^2^) were cultured with 2 % FBS and 10 μM all-trans retinoic acid at bottom of the well for one week (Fig. 1b, e). As well, for other triple co-culture models, mixed glial cells were seeded on a 24-well plate with a concentration of 4×10^4^ cells/cm^2^ and 10 % FBS (Fig. 1c, e).

In co-culture models with astrocytes (Fig. 1e), in a first step, 1321N1 cells were seeded on the underside of the inserts with 1.5×10^4^ cells/cm^2^ concentration and incubated for 2-3 h inversely, followed by 24 h in media to compatibility. After 24 h, hCMEC/D3 cells were seeded on the apical side, and after formation of a monolayer, inserts were transferred to wells containing differentiated WT SH-SY5Y or O-αSN SH-SY5Y cells, as well as primary mixed glial cells. Primary mixed glial cells were treated with WT SH-SY5Y or O-αSN SH-SY5Y cells conditioned media as well as LPS (10 μg/mL) or 12-h αSN-AGs (5% v/v). WT SH-SY5Y cells were treated with 12-h αSN-AGs (5% v/v). TEER was measured after formation of monolayer (0 h) and before transferring which considered as 100 %. The integrity was then assessed by TEER 24 and 48 h after transfer. Triple co-culture models and their treatments, as well as experimental design, are detailed in Fig. 1b, c, e and **Table** 2.

**Table 2.** Overview of co-culture groups and their treatments for measuring TEER

| Groups | Cell plated on apical &<br>basolateral side of the insert | Cell plated on<br>lower compartment | Treatment of lower compartment |
| --- | --- | --- | --- |
| **Group 1** | hCMEC/D3 & 1321N1 | WT SH-SY5Y | --- |
| **Group 2** | hCMEC/D3 & 1321N1 | WT SH-SY5Y | Exogenous 12-h αSN-AGs |
| **Group 3** | hCMEC/D3 & 1321N1 | O-αSN SH-SY5Y | --- |
| **Group 4** | hCMEC/D3 & 1321N1 | Primary mixed glia | WT SH-SY5Y conditioned media |
| **Group 5** | hCMEC/D3 & 1321N1 | Primary mixed glia | O-αSN SH-SY5Y conditioned media |
| **Group 6** | hCMEC/D3 & 1321N1 | Primary mixed glia | Exogenous 12-h αSN-AGs |
| **Group 7** | hCMEC/D3 & 1321N1 | Primary mixed glia | LPS |

Exogenous 12-h αSN-AGs α-synuclein incubated for 12 h under fibrillation condition, WT SH-SY5Y Wild type SH-SY5Y cells, O-αSN SH-SY5Y Tet-Off SH-SY5Y cells expressing αSN in the absence of doxycycline

#### Measurement of NO production

The amount of NO was estimated by measuring the nitrite concentration, a stable metabolite of NO, in the supernatant of the primary mixed glial or hCMEC/D3 cell cultures using the Griess Reaction assay.

For this purpose, primary mixed glial or hCMEC/D3 cells were seeded on 24-well plates and 96-well plates with 4×10^4^ cells/cm^2^ and 1.5×10^4^ cells/cm^2^ density, respectively. After 24 h, hCMEC/D3 cells were treated with 12-h αSN-AGs or 24-h αSN-AGs (10% v/v). After 7 days, the primary mixed glial cells were treated with LPS (10 μg/mL), O-αSN SH-SY5Y conditioned media (under the conditions for inducing αSN expression, see previous sections), or 12-h αSN-AGs (5% v/v). After treatment time (6 and 24 h for hCMEC/D3 cells and 48 h for primary mixed glial cells), the Griess Reaction assay, which is based on the two-step diazotization reaction, was performed. To increase the sensitivity of the Griess reaction, the reagents were added in two steps and at 4 °C according to previous study [44]. Firstly, sulphanilamide (1% (w/v) in 5% H_3_PO_4_) was added to the supernatant of the cultures for 10 min, after which NED (N-1-naphthylethylenediamine dihydrochloride (0.1% w/v) was added and the solution incubated for another 10 min. The absorbance of the samples was measured at 540 nm using a Biotek Epoch 2 microplate spectrophotometer (Agilent, Santa Clara, CA). The standard curve of sodium nitrite (0-100 μM) was used to determine the NO concentration. The details of the studied groups are indicated in **Table 3**.

**Table 3.**
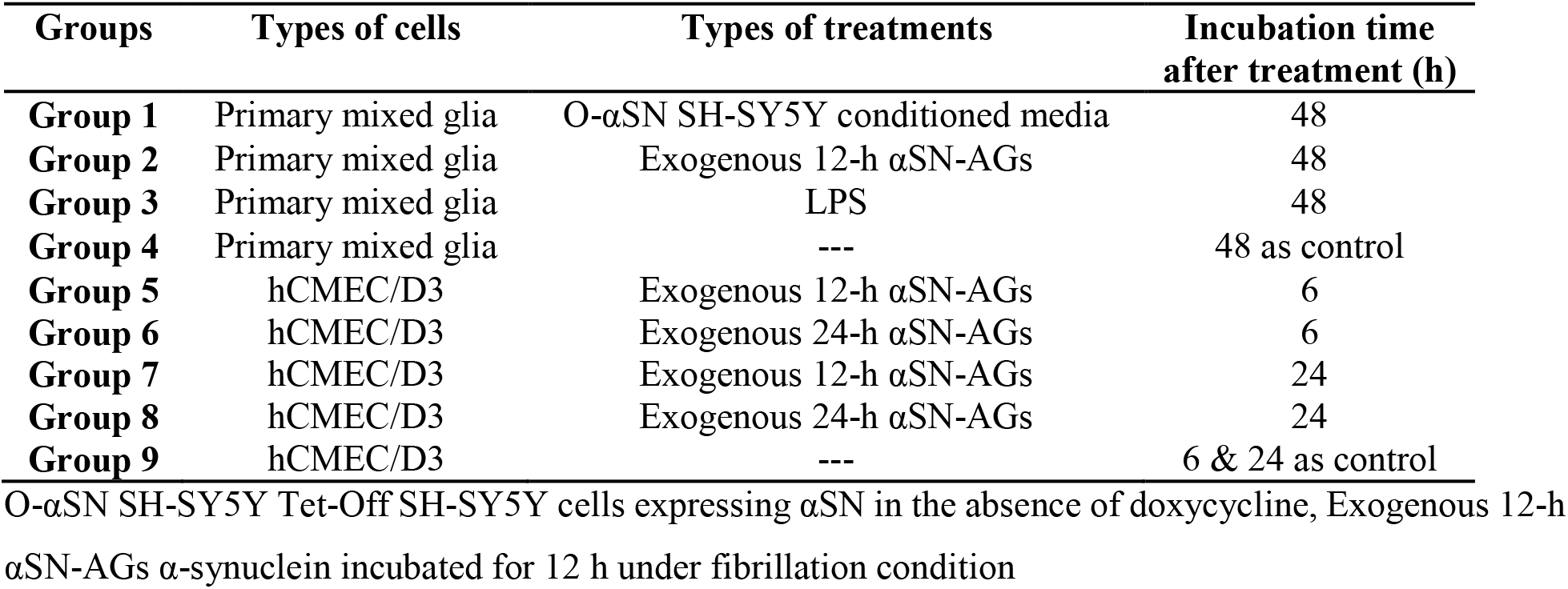
Overview of treatment groups for assessment of NO production

#### Exploring the transmigration of Jurkat cells across the BBB

Transmigration assay across the BBB was performed based on the previous description [45].

#### Experimental design for measuring transmigrated Jurkat cells across the BBB

hCMEC/D3 cells were seeded on the inserts with 8.0 μm pore size as mentioned previously. As shown in Fig. 1d, f, after formation of the monolayer, the inserts were transferred to a new plate where each well had different conditions, including: (group 1) simple media as control, (group 2) media with 5 % v/v of 12-h αSN-AGs, (group 3) culturing O-αSN SH-SY5Y cells on the bottom of wells for a week in differentiating and αSN expressing conditions, (groups 4,5, respectively) culturing primary mixed glia cells in the vicinity of 12-h αSN-AGs (group 4) or O-αSN SH-SY5Y conditioned media (cultured under conditions to induce αSN expression, see previous sections) (group 5). After 16 h, Jurkat cells were added on the apical side of the inserts, which had already been cultured with hCMEC/D3 monolayer. After 8 h, the migrated Jurkat cells were collected from the lower compartment, centrifuged and re-suspended in 200 μL media, and the numbers of the collected cells were counted using a Neubauer hemocytometer. The rate of the migration was normalized to the input cells. **Table 4** and Fig. 1d, f, describe co-culture cells and their treatments, as well as the experimental design.

**Table 4.** Overview of co-culture groups and their treatments for transmigration assay

| **Groups** | **Cell plated on apical side of the insert** | **Cell plated on lower compartment** | **Treatment of lower compartment** |
| --- | --- | --- | --- |
| **Group 1** | hCMEC/D3 | --- | --- |
| **Group 2** | hCMEC/D3 | --- | Exogenous 12-h αSN-AGs |
| **Group 3** | hCMEC/D3 | O-αSN SH-SY5Y | --- |
| **Group 4** | hCMEC/D3 | Primary mixed glia | O-αSN SH-SY5Y conditioned media |
| **Group 5** | hCMEC/D3 | Primary mixed glia | Exogenous 12-h αSN-AGs |

Exogenous 12-h αSN-AGs α-synuclein incubated for 12 h under fibrillation condition, O-αSN SH-SY5Y
Tet-Off SH-SY5Y cells expressing αSN in the absence of doxycycline

#### Measuring the extent of interaction of different αSN conformers with cells

αSN was first conjugated with FITC, purified and then added to different cells.

#### Labeling of the αSN with FITC

αSN was fibrillated as mentioned above. Samples were collected at different times and incubated with FITC solution (dissolved in DMSO) at a final concentration of 1400 μM FITC/ 70 μM αSN under continuous stirring for 3 h at 4 °C. Unreacted FITC was removed by size exclusion chromatography using a PD-10 column while collecting fractions containing labeled αSNs [46]. Protein concentration and moles of FITC per mole of protein (F/P) were calculated as follows [47, 48]:

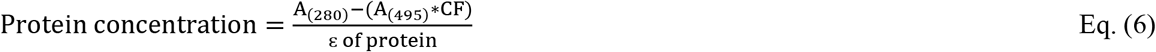

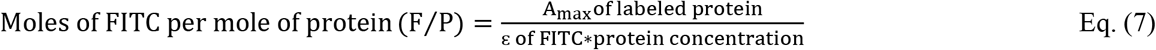

where ɛ represents the molar extinction coefficient factor and CF the correction factor. The fluorescence intensity was assayed at 495 nm and 280 nm (excitation slit width 5 nm) and 518.95 nm and 310 nm (emission slit width 10 nm) for FITC and αSN, respectively [49].

#### Detection of FITC-αSN-AGs in hCMEC/D3, HUVEC and 1321N1

hCMEC/D3, HUVEC and 1321N1 cells were seeded on 24-well plates with 10^5^ cells/cm^2^ density, and after 24 h, treated with FITC-αSN-AGs at a final concentration of 10% v/v. After 3 h, cells were trypsinized and re-suspended in PBS, after which fluorescence intensity was measured with excitation at 495 nm and emission spectra in the range of 505 to 550 nm. The excitation and emission slit widths were 10 nm for HUVEC cells and 10 and 20 nm for hCMEC/D3 and 1321N1 cells.

#### Statistical analysis

All experiment results were expressed as mean±SD, and each experiment was repeated in triplicate, unless otherwise indicated. The statistically significant differences between groups were determined by one-way ANOVA (Tukey’s Honest Significant Difference post-hoc) with a conventional threshold of p ≤ 0.05 using SPSS software v.16.0.

## Results

### Production and validation of αSN-AGs during fibrillation

To prepare fibrils of αSN, we incubated the protein under conditions inducing amyloid fibrillation and analyzed samples after 3, 7, 12, and 24 h using ThT fluorescence intensity and CR absorbance (Fig. S1a, b), supplemented with AFM and ThT fluorescence microscopy images at 3, 12, and 24 h (Fig. S2a, b, c). All data confirmed that αSN formed aggregates (αSN-AGs) consisting mainly of a mixture of fibrils of different lengths interspersed with smaller species which are likely oligomers (particularly prevalent around 3 and 12 h).

### αSN amyloid aggregates have detrimental effects on endothelial cells

To explore the effects of differently aged αSN-AGs on hCMEC/D3 cells, we measured cell viability (MTT assay and DAPI staining), mitochondrial membrane potential (ΔΨm) and intracellular ROS/NO production.

#### Cell viability

Cells were treated with fresh αSN and different aggregated species of αSN (3-, 7-, 12-, and 24-h αSN-AGs). The MTT assay showed that the viability of hCMEC/D3 cells diminished significantly, especially when treated with 7-h and 12-h αSN-AGs (Fig. 2a) with a clear dose-response effect (Fig. 2b). DAPI staining confirmed cellular toxicity with increased chromatin condensation and nuclear fragmentation after treating with 12-h or 24-h αSN-AGs (Fig. S3a, b).

**Fig. 2.**
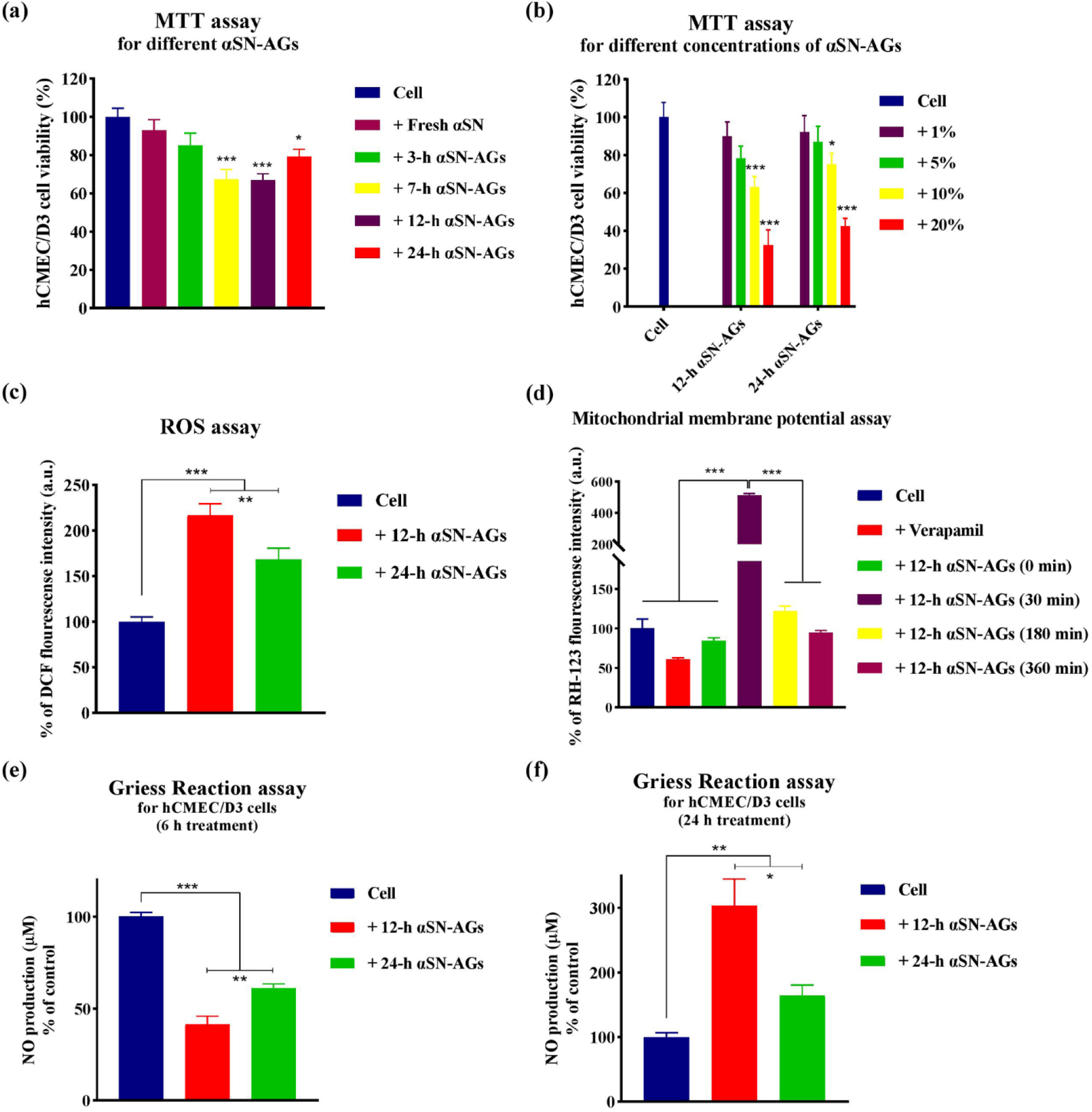
Exploring the effects of αSN-AGs on BBB endothelial cells. The effects of αSN-AGs were evaluated by assessing cell viability, mitochondrial membrane potential, intracellular ROS activity, and NO production. The viability of hCMEC/D3 cells in the presence of (a) different αSN-AGs (10% v/v), or (b) different concentrations of 12-h and 24-h αSN-AGs as measured by MTT. (c) Effect of 12-h and 24-h αSN-AGs (10% v/v) on the hCMEC/D3 intracellular ROS content. The DCF emissions of samples treated with αSN-AGs were normalized to that of control. (d) Mitochondrial membrane potential assay of hCMEC/D3 cells in the absence and presence of 12-h αSN-AGs (10% v/v) using RH-123. (e, f) Measurement of NO levels in the supernatant of hCMEC/D3 cell culture in the absence and the presence of 12-h and 24-h αSN-AGs (10% v/v) for 6 h and 24 h, by Griess Reaction assay. * indicates significant differences between treated and control samples, as well as between marked groups. (Mean±SD, * P value ≤ 0.05, ** P value ≤ 0.01, *** P value ≤ 0.001).

#### ROS production

Similarly, the DCFH-DA assay revealed that cells treated with 12-h or 24-h αSN-AGs showed significant increases in their cytoplasmic ROS content (Fig. 2c).

#### Mitochondrial membrane potential

ΔΨm was reduced by αSN-AGs as ascertained using the fluorescent dye RH-123. In the healthy state, RH-123 accumulates in active and intact mitochondria with self-quenching as determined by adding 50 μM verapamil (which as a P-gp inhibitor will accelerate uptake of RH-123 into hCMEC/D3 cells); this is confirmed by the very modest effect of 12-h αSN-AGs at time zero when it has not had significant time to have an effect. However, after 30 min of exposure to 12-h αSN-AGs, the signal has increased substantially, possibly due to αSN-AGs impacts on the mitochondrial membrane potential and entrance of RH-123 into the cytoplasm and un-quenching. However, the intensity was reduced after 180 and 360 min, probably due to RH-123’s inability to enter damaged mitochondria (more loss of ΔΨm) (Fig. 2d).

#### Production of NO and NO-derived oxidants

Cells were treated with 12-h αSN-AGs and 24-h αSN-AGs for 6 and 24 h. After 6 h, the NO content significantly declined in 12-h αSN-AGs and 24-h αSN-AGs compared to the control group (Fig. 2e). In contrast, 12-h αSN-AGs and 24-h αSN-AGs increased NO levels in the supernatant compared to the control group when exposed to hCMEC/D3 cells for 24 h (Fig. 2f). This may be due to differences in expression levels of two isoforms of nitric oxide synthase enzyme (eNOS and iNOS), through which the elevation of NO may be related to iNOS activation over time (see Discussion).

### αSN-AGs impair BBB functionality

TEER and impermeability are two central parameters to evaluate the functionality of the BBB. To determine the effects of αSN-AGs on the TEER and permeability values, hCMEC/D3 cells were cultured on the apical side of transwell tissue culture inserts (Fig. 1a). Formation of the cell monolayer was monitored by phase contrast microscopy during the days of experiment (Fig. 3a).

**Fig. 3.**
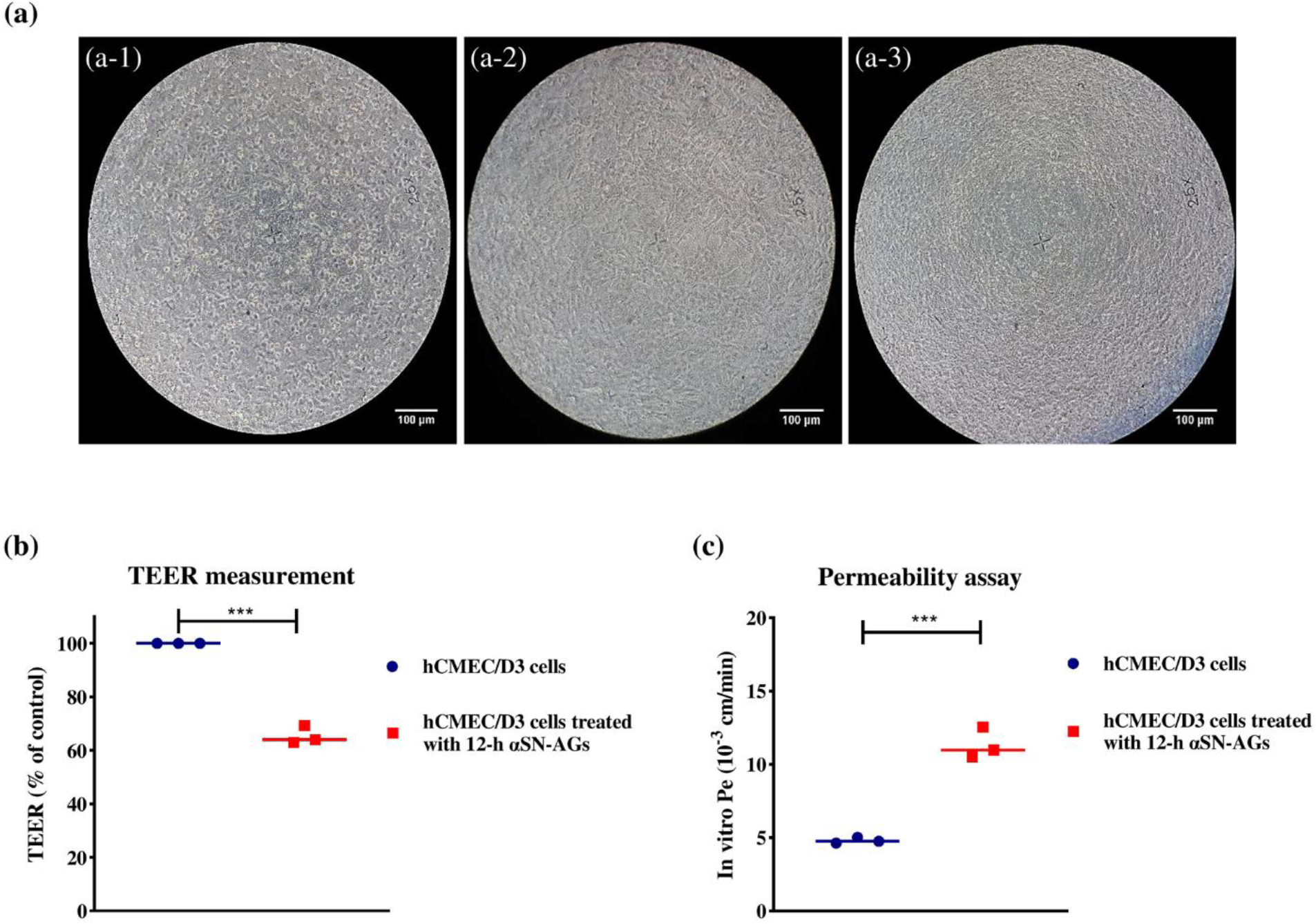
Integrity assessment of the monolayer in the absence of αSN using TEER and permeability measurements. hCMEC/D3 cells formed a monolayer in the apical side of insert of transwell (images are taken on the first day after culturing (a-1), the day in which the monolayer has not yet been completely formed (a-2), and the day that monolayer formed completely (a-3)). The cells were then treated with 12-h αSN-AGs (5% v/v). TEER measurement (b) and permeability assay (c) showed significant differences between the treated and control samples. TEER and permeability values were measured and determined using Eqs. 3–5. * indicates significant differences between treated and untreated samples, (Mean±SD, *** P value ≤ 0.001).

We measured BBB characteristics when treated with αSN. For this purpose, after formation of a monolayer, the cells were treated with 12-h αSN-AGs on the apical side, and then TEER and permeability values were measured. When cells were treated with 12-h αSN-AGs, the TEER values decreased significantly while the permeability values increased (Fig. 3b, c).

### αSN-AGs disrupted the wound healing performing of brain microvascular endothelial cells

Cellular migration of hCMEC/D3 cells was assessed by wound healing assay. As shown in Fig. 4a, untreated cells (group 1), expressed a sheet or grouping migration and stayed connected to each other. Cells treated with 12-h αSN-AGs or 24-h αSN-AGs (groups 2 and 3, respectively) showed individual cell migration (particularly in group 2), which is not an appropriate response for wound healing and angiogenesis (Fig. 4b, c). The wound healing assay was also done for 1321N1 and HUVEC cells as representatives of non-endothelial cells and non-BBB endothelial cells, respectively. 1321N1 cells also showed single cell movement without any angiogenic behavior (Fig. S4a). Interestingly, HUVEC cells underwent collective cell movement and filled the open area to some extent without single cell movement, similar to the untreated control group of hCMEC/D3 cells (Fig. S4b). As shown in Fig. 4b, the control group of hCMEC/D3 cells and HUVEC cells showed a similar process of filling the gap. In addition, treated hCMEC/D3 cells behaved almost the same as 1321N1 cells with non-endothelial features. Untreated hCMEC/D3 cells (group 1) showed the highest percentage of wound closure but this dropped significantly when cells were treated with 12-h αSN-AGs or 24-h αSN-AGs (Fig. 4c).

**Fig. 4.**
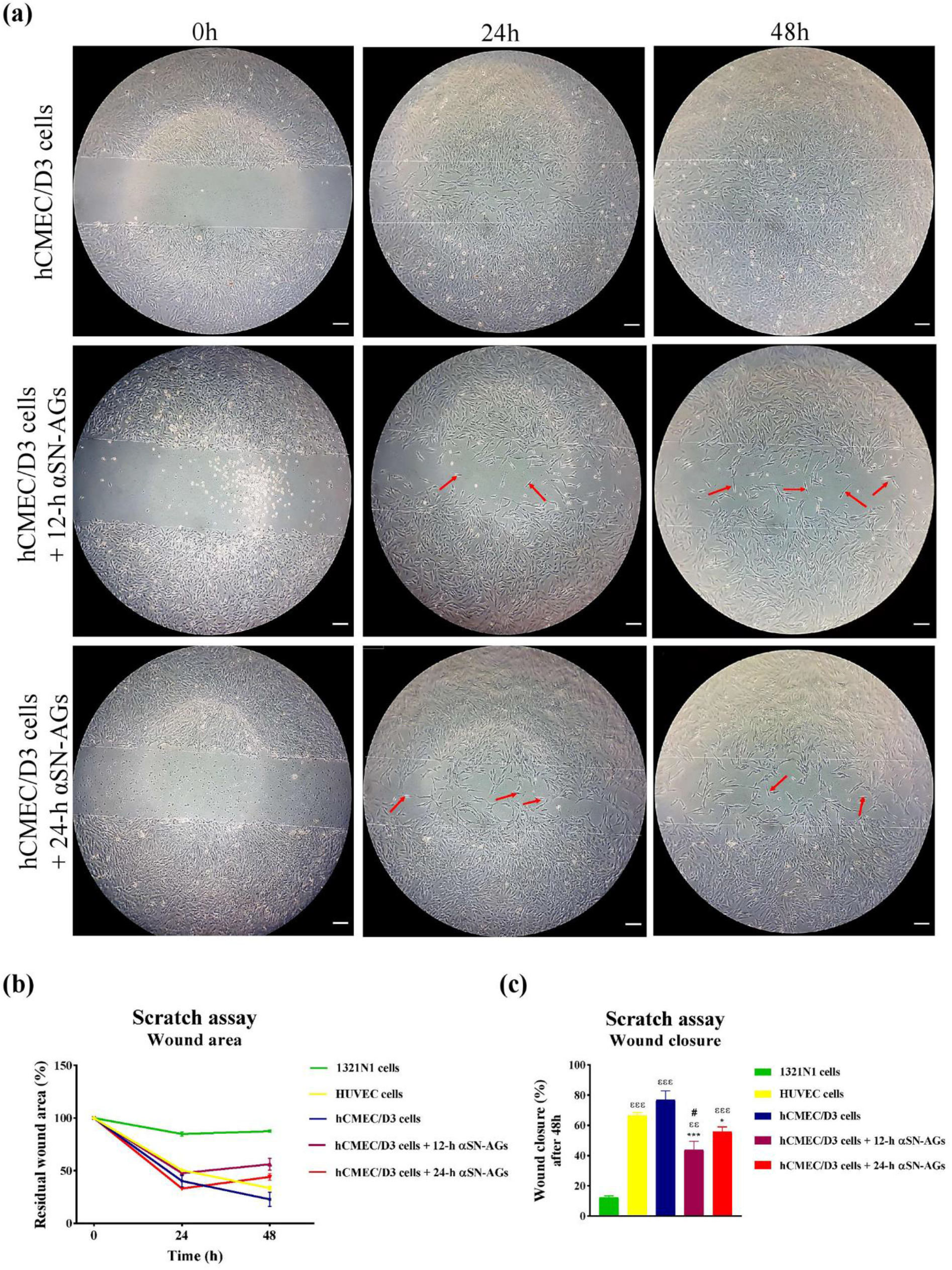
*In vitro* wound healing assay (scratch assay). hCMEC/D3 cells were grown to full confluent density, scratched and then treated with 12-h and 24-h αSN-AGs (5% v/v). The scratched field area and scratched field closure were calculated using ImageJ and Eqs. (1) and (2). (a) Cell morphology of hCMEC/D3 cells in the absence and presence of αSN-AGs. Isolated cells with random movements were shown by red arrows. Quantitative analysis of residual scratched area (b) and wound closure rate after 48 h (c). Quantification of scratched area and wound closure were calculated using ImageJ software according to Eqs. (1) and (2). * indicates significant differences between hCMEC/D3 cells and those treated with αSN-AGs. # indicates significant differences between HUVEC cells and hCMEC/D3 cells treated with αSN-AGs. ɛ indicates differences between 1321N1 cells and other groups. (Mean±SD, *, # P value ≤ 0.05, ɛɛ P value ≤ 0.01, ***, ɛɛɛ P value ≤ 0.001). Scale bar: 50 μm.

### αSN-AGs could stimulate the inflammatory response of glial cells either in exogenous or in endogenous overexpressed forms

To identify the effects of αSN on the activation of glial cells and inflammation responses, primary mixed glial cells were treated with 1) the conditioned media derived from O-αSN SH-SY5Y cells, 2) 12-h αSN-AGs, 3) LPS, and 4) no treatment (see **Table 3** for details). All three treated groups showed elevated NO levels which represented the activation of primary mixed glial cells (Fig. 5a). Activation of glial cells was confirmed by morphological analysis [50]. Microglial cells in resting or non-activated state have a comparatively small cell body with highly ramified processes (outgrowths). However, in the activated state when exposed to LPS, αSN-AGs or O-αSN SH-SY5Y-derived conditioned media, microglial cells undergo hypertrophy and retract their processes. All treated groups showed microglia with thicker primary processes and retracting secondary processes in first stage of activation (mildly activated hyper ramified microglia), and un-ramified (activated) microglia (in the last stages of activation) with bushy morphology, swollen shape and enlarged cell body and truncated processes, and amoeboid forms with rounded macrophage-like morphology (Fig. 5b).

**Fig. 5.**
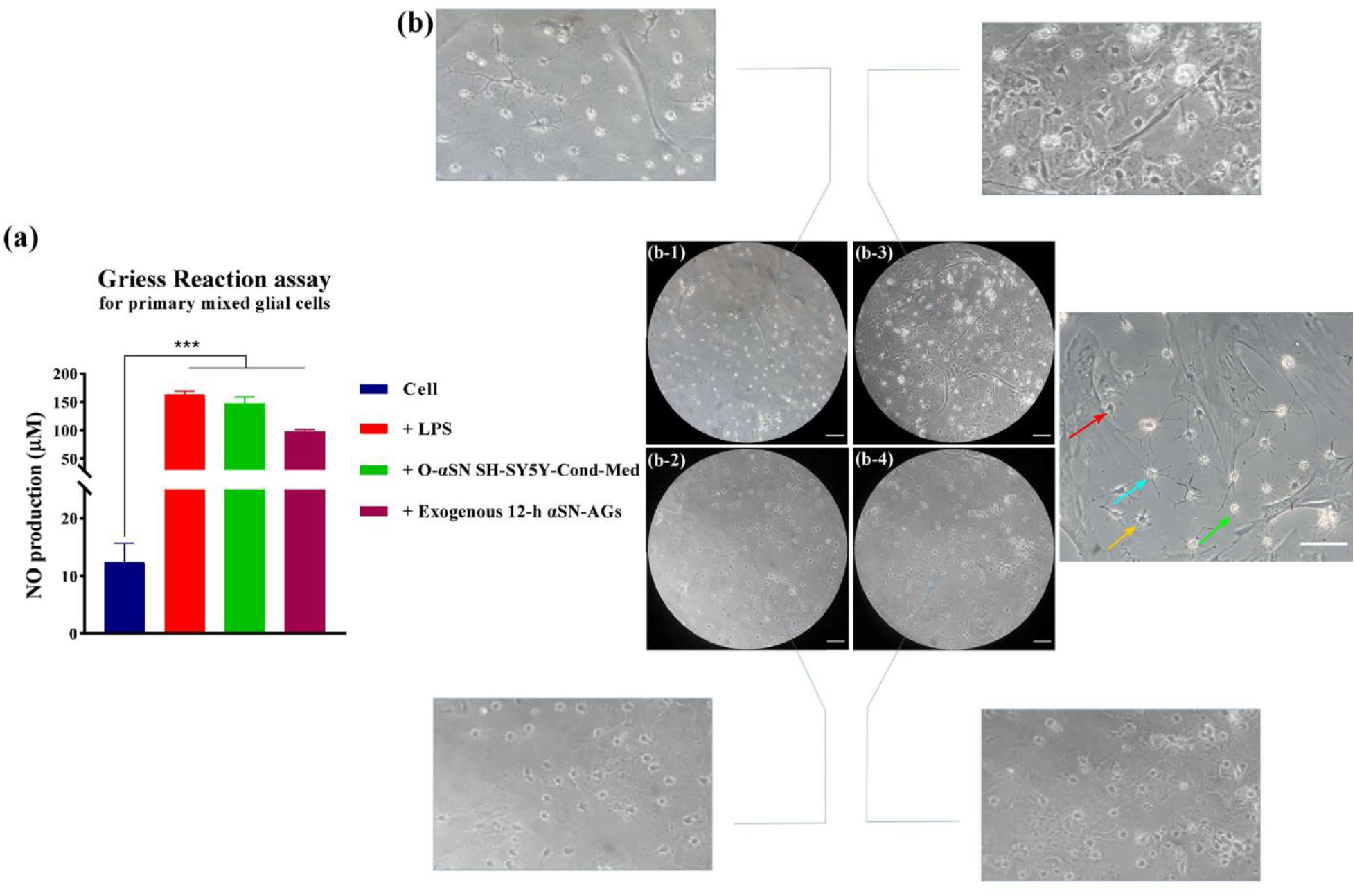
Inflammatory activation assay. (a) NO production was assessed in the supernatants of primary mixed glial cell cultures after 48 h treatment using Griess Reaction assay. (b) Morphological analysis of the microglial cells in (b-1) no treating condition with a dominant morphology in the resting state characterized by microscopic or small cell bodies and thin branching processes with ramified appearance, (b-2) treated with LPS (10 μg/mL) with a dominant morphology of activated state characterized by bushy and amoeboid morphology (un-ramified), (b-3) treated with 12-h αSN-AGs (5% v/v) represents a mixture of hyper ramified and un-ramified (activated), and (b-4) treated with O-αSN SH-SY5Y-derived conditioned media display un-ramified (activated). Different activation stages of microglia are indicated by arrows; green: resting microglia, cyan: mildly activated hyper ramified, orange: hyper-ramified microglia and red: amoeboid microglia. * indicates significant differences between treated samples and that of control (Mean±SD, *** P value ≤ 0.001). Scale bar: 50 μm.

### Various NVU cells or their conditioned media could alter impacts of αSN on BBB

The endothelial cells are near neurovascular unit (NVU) cells. In this regard, the protective effects of conditioned media derived from astrocytes on neurons and vice versa against three different neurotoxicants have been reported [51]. This was carried out using transwell non-contact co-culture systems (plated on apical side of insert and bottom of the well) [51]. Accordingly, the effects of conditioned media derived from astrocytes and dopaminergic neurons (Ast-Cond-Med and Neu-Cond-Med) in the absence and presence of αSN-AGs were evaluated on hCMEC/D3 cells (Fig. 6). As expected, 12-h αSN-AGs reduced endothelial cell viability by 60% in the comparison with untreated cells, while addition of conditioned media from both types of cells reduced the cytotoxicity of αSN-AGs significantly (Fig. 6a, b). We also measured the cell viability of these cells after treating them with αSN-AGs which reduced the viability of 1321N1 and SH-SY5Y to approximately 87 % and 75 %, respectively (Fig. S5).

**Fig. 6.**
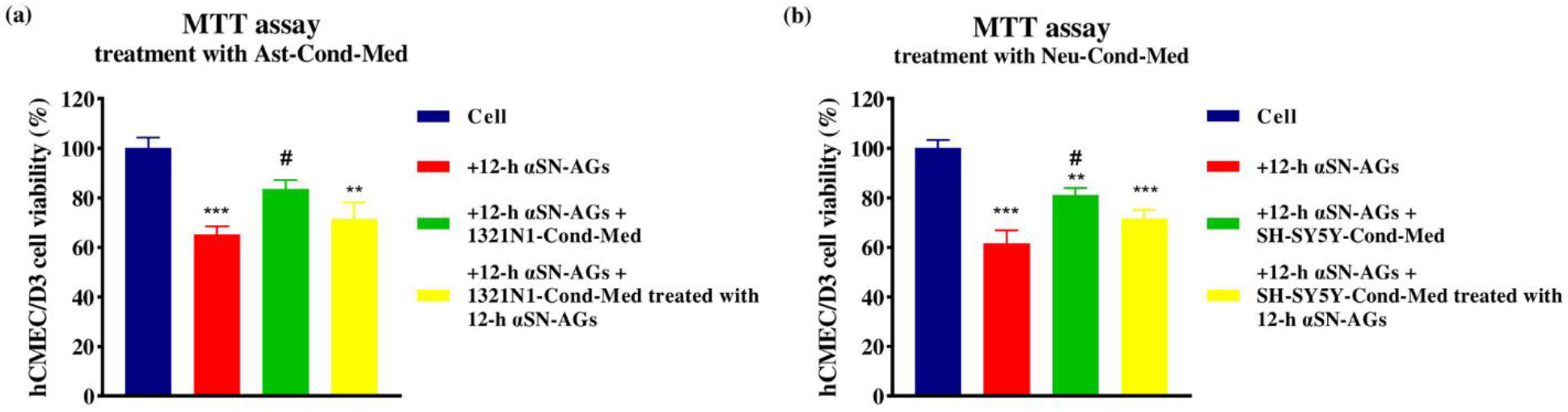
Assessment of the effect of Ast-Cond-Med and Neu-Cond-Med on the toxicity of αSN-AGs. The viability of hCMEC/D3 cells in the presence of 12-h αSN-AGs (10% v/v) and conditioned media derived from (a) 1321N1 and (b) SH-SY5Y treated without or with (5% v/v) 12-h αSN-AGs, measured by MTT assay. * indicates statistically significant differences between treated and control samples; # indicates statistically significant differences between hCMEC/D3 cells treated with αSN-AGs and those treated with free Ast-Cond-Med and Neu-Cond-Med. (Mean±SD, *, # P value ≤ 0.05, ** P value ≤ 0.01, *** P value ≤ 0.001).

We also evaluated the responses of the hCMEC/D3 cells to αSN-AGs when co-cultured with astrocytes and neurons. For this purpose, 1321N1 cells were initially seeded on the basolateral side of the insert, and after 24 h, hCMEC/D3 cells were seeded on the apical side. Then, inserts were transferred to a plate cultured with SH-SY5Y cells (wild or αSN overexpressing types), which were seeded before transferring (Fig. 1b, e). In parallel, one group of WT SH-SY5Y cells were treated with 12-h αSN-AGs (Fig. 1b, e). To address the effect of including microglia and inflammatory responses, TEER values were measured in 4 treatment groups containing hCMEC/D3 and 1321N1 cells on the apical and basolateral sides of the insert, respectively, transferred to a plate cultured with: 1) primary mixed glial cells with WT SH-SY5Y cells conditioned media (healthy control), 2) primary mixed glial cells treated with O-αSN SH-SY5Y cells conditioned media (one week without doxycycline) (pathological condition inducing inflammatory response), 3) primary mixed glial cells treated with LPS (10 μg/μL) (positive control inducing inflammatory response), and 4) primary mixed glial cells treated with 12-h αSN-AGs (pathological condition inducing inflammatory response), as shown in Fig. 1c, e. TEER values were assessed for up to 48 h.

As shown in Fig. 7a, monolayers of hCMEC/D3 and 1321N1 were formed on the apical and basolateral sides of inserts, respectively, and were imaged by phase contrast microscopy. The left-side image shows the monolayer of hCMEC/D3 cells on the apical side, while the astrocyte monolayer could be imaged on the basolateral side (right-side image) by scratching on the apical side of one insert. Under all conditions, endogenous or exogenous αSN could be expected to affect TEER values. In co-culture with SH-SY5Y cells, exogenous αSN and endogenous αSN overexpressed in O-αSN SH-SY5Y cells reduced the TEER mildly compared with WT SH-SY5Y cells, however was not significant (Fig. 7b). Moreover, as shown in Fig. 7c, in the inflammatory and non-inflammatory conditions in co-culture of primary mixed glial cells, exogenous αSN had essentially the same effect as LPS and caused a significant reduction in TEER values. Similarly, endogenous αSN in co-culturing of primary mixed glial cells in the presence of O-αSN SH-SY5Y cells conditioned media (O-αSN SH-SY5Y-Cond-Med) generated inflammatory status and decreased TEER value.

**Fig. 7.**
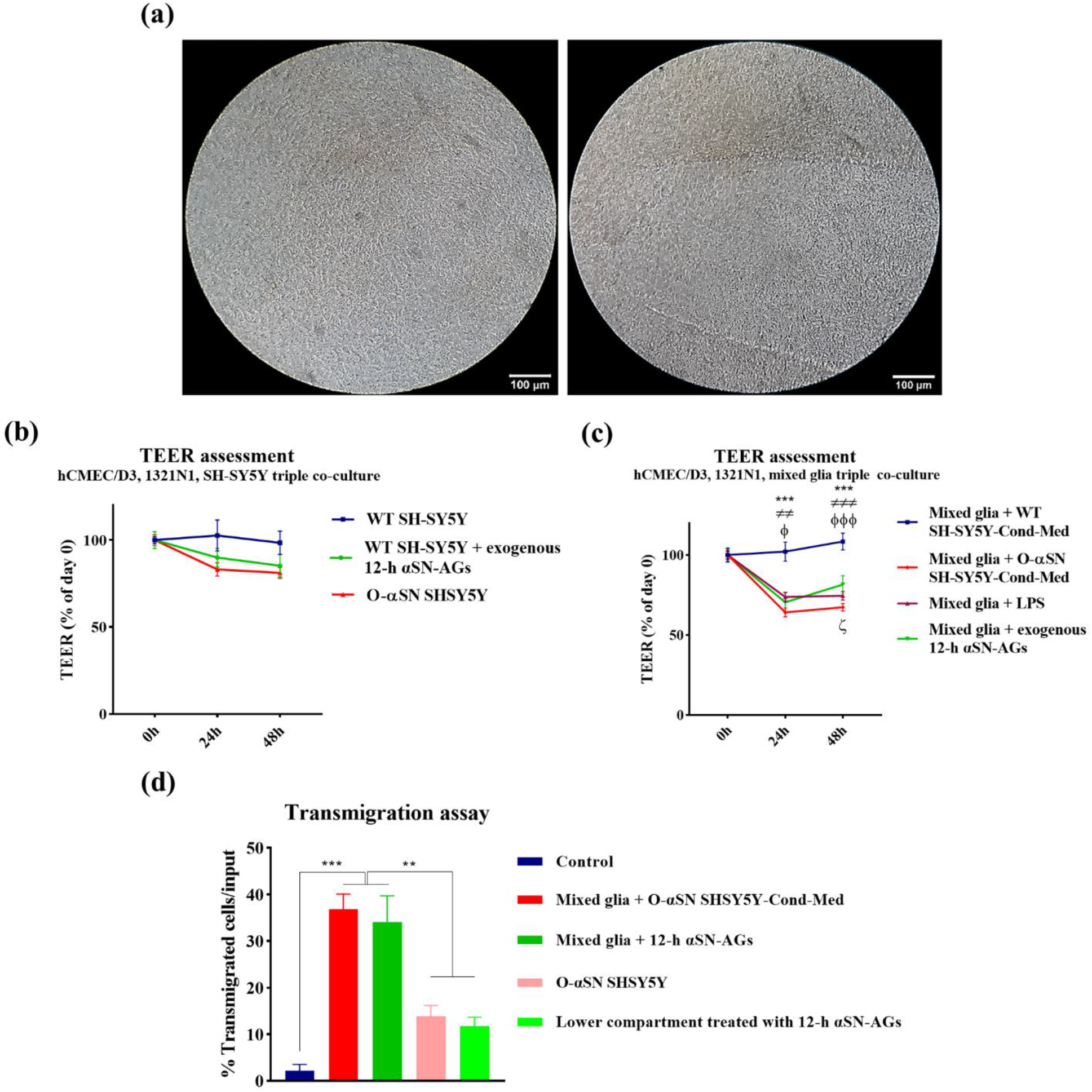
Assessment of the integrity of the BBB model in different conditions and co-cultures scenario. The integrity was assessed by measuring TEER parameter and evaluation of T cell migration across BBB. hCMEC/D3 and 1321N1 cells were cultured in the apical and basolateral sides of insert of transwell, respectively. To ascertain that 1321N1 cells were settled in the basolateral side of insert together with hCMEC/D3 on the apical side, we scratched the apical side (a right-side image). The inserts were then transferred to plates containing WT SH-SY5Y, WT SH-SY5Y treated with 12-h αSN-AGs (5% v/v) or O-αSN SH-SY5Y that has already been cultured and differentiated for a week, and TEER was measured up to 48 h (b). The same procedure was carried out with a 24-well plate containing mixed glia treated with 12-h αSN-AGs (5% v/v), LPS (10 μg/μL), WT SH-SY5Y -derived conditioned media (WT SH-SY5Y-Cond-Med) or O-αSN SH-SY5Y -derived conditioned media (O-αSN SH-SY5Y-Cond-Med) and TEER was measured up to 48 h (c). *, ǂ and φ indicate significant differences between WT SH-SY5Y-Cond-Med and O-αSN SH-SY5Y-Cond-Med, 12-h αSN-AGs and LPS treatment, respectively. As well, ζ indicates significant differences between O-αSN SH-SY5Y-Cond-Med and 12-h αSN-AGs treatment. (Mean±SD, *, ǂ, φ, ζ P value ≤ 0.05, **, ǂǂ P value ≤ 0.01, ***, ǂǂǂ, φφφ P value ≤ 0.001). Transmigration assay (d), after monolayer formation of hCMEC/D3 cells in the apical side of the insert, transferred to 24-well plate containing untreated media, media treated with 12-h αSN-AGs or wells containing O-αSN SH-SY5Y cells or mixed glia treated with 12-h αSN-AGs or O-αSN SH-SY5Y conditioned media. Then Jurkat cells were added to the apical side of insert and their migrations rate through hCMEC/D3 monolayer were assessed. (Mean±SD, * P value ≤ 0.05, ** P value ≤ 0.01, *** P value ≤ 0.001).

### In the vicinity of αSN, the penetration of T cells through BBB accelerated, particularly under inflammatory stimulation

To identify the effect of αSN-AGs on T cell recruitment to CNS in the pathological conditions, we explored the migration of Jurkat cells across the BBB endothelial monolayer. For this, hCMEC/D3 cells after monolayer formation (confirmed by TEER) were treated with various preparations in the lower compartment (**Table 4**): (1) untreated media as control group, (2) treated media with 12-h αSN-AGs, and other groups plated with (3) O-αSN SH-SY5Y cells and (4, 5) mixed glia that were treated with (4) 12-h αSN-AGs or (5) O-αSN SH-SY5Y conditioned media (Fig. 1d, f). Migration of Jurkat cells into the lower chamber after 8 h significantly increased in co-culture groups with primary mixed glia and inflammatory conditions compared with non-inflammatory conditions (Fig. 7d).

### Diversity in the interactions of αSN-AGs with BBB endothelial cells

To probe the overall interaction of αSN with hCMEC/D3, αSN was labeled with FITC after incubation for different periods under fibrillation conditions and the labelling was quantified by absorption. As seen in Fig. 8a and Table S1, the extent of labeling declines with ageing of αSN under fibrillation conditions and is essentially abolished after 24 h. After labeling of αSN (fresh αSN and 3-, 7-, 12-, and 24-h αSN-AGs), FITC-αSN was added to the culture of hCMEC/D3, HUVEC and 1321N1 cells. As depicted in Fig. 8b, hCMEC/D3 showed more FITC fluorescence intensity when incubated with 7-h, 12-h and 24-h αSN-AGs in comparison with other αSN-AGs. We also determined that in another kind of endothelial cell, HUVEC, the fluorescence intensity was lower in comparison to BBB endothelial cells. In addition, with the same manner, 1321N1 cells have a higher tendency than endothelial cells to interact with αSN-AGs, predominantly 12-h αSN-AGs and 24-h αSN-AGs.

**Fig. 8.**
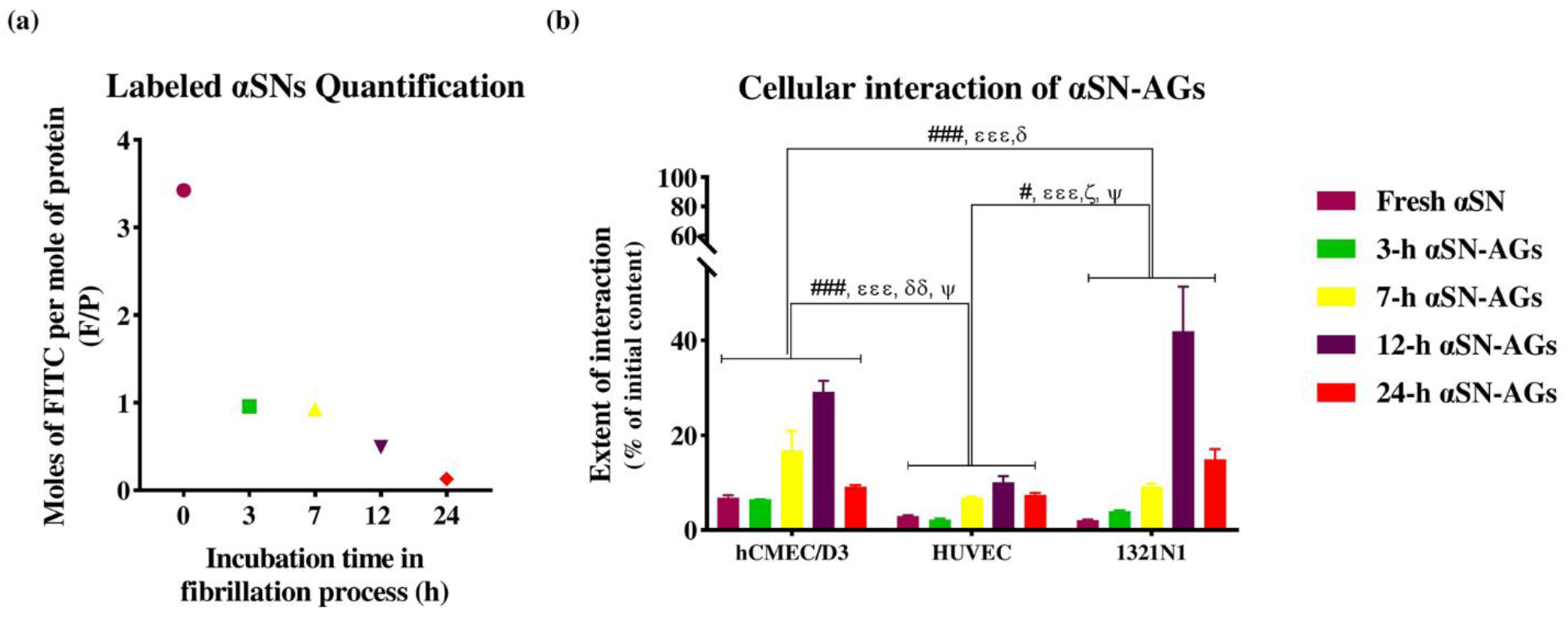
Exploring the interaction of αSN with BBB endothelial cells using labeled αSN. (a) Moles of FITC per mole of αSN (F/P) as a function of the extent of fibrillation, calculated according to Eqs. (6) and (7). αSN-AGs were obtained by incubation under fibrillation conditions for different periods of time. (b) FITC fluorescence intensity was measured to analyze the extent of interaction between different conformers of αSN and hCMEC/D3, HUVEC and 1321N1 cells. #, ɛ, δ, ζ, and Ψ indicate FITC labeled- αSN 0-h αSN-AGs (fresh αSN), 3-h αSN-AGs, 7-h αSN-AGs, 12-h αSN-AGs, 24-h αSN-AGs respectively, which the significance between groups is indicated in the diagram separately. (Mean±SD, #, ɛ, δ, ζ, Ψ P value ≤ 0.05, ##, ɛɛ, δδ, ζζ, ΨΨ P value ≤ 0.01, ###, ɛɛɛ, δδδ, ζζζ, ΨΨΨ P value ≤ 0.001).

## Discussion

The BBB is a complex organ which constitutes a highly regulated interface between the CNS and peripheral circulation. It plays a particular role in controlling the passage of substances and immune cells between the brain and outside of it [1]. This role is largely due to its specified endothelial cells and their interaction with other cells of NVU, including microglia and neuronal cells [1]. There is widespread agreement that BBB plays a critical role in the progression and severity of brain diseases, particularly NDs [52].

Motivated by the fact that αSN aggregation in patients with NDs, especially PD, is observed both within and outside the brain [19], as well as its paramount role in the pathogenesis of synucleinopathies and the association of BBB disruption in the progression of the diseases [5–7], we studied αSN effects on *in vitro* models of BBB and its function from both apical and basolateral sides along with BBB related cells.

### αSN-AGs showed direct destructive impacts on BBB endothelial cells

First of all, treatment of BBB endothelial cells with αSN pre-incubated under fibrillation conditions caused a significant decrease in cell viability in a time- and concentration-dependent manner. In addition, the intra-cellular ROS and NO content changed considerably when the cells were treated with αSN-AGs. The role of NO in endothelial cells is somewhat unclear. NO produced by inducible nitric oxide synthase (iNOS) can induce apoptosis in vascular endothelial cells [53], while NO produced by endothelial nitric oxide synthase (eNOS) protects against excessive ROS and also endothelial cells apoptosis [54]. Note that iNOS expresses later than eNOS [55], which suggests that in our study, NO content could be elevated over time through iNOS activation with more hazardous effects, as observed in our results after 24 h.

NO derivatives and NO redox forms (NO^*^, NO^+^, NO^−^) are involved in BBB dysfunction and BBB permeability from mild to severe damage [56]. Moreover, matrix metalloproteinases (MMPs) are implicated in BBB disruption and can be activated by NO and its derivatives [57].

Treatment with αSN-AGs led to mitochondrial dysfunction, in accordance with other studies [58, 59]. Specifically, treatment with αSN led to reduced mitochondrial membrane potential and increased intracellular ROS. Mitochondria play crucial roles in BBB endothelial cells integrity by maintaining ion homeostasis between blood and brain, as well as bolstering its impermeability [60]. As a consequence, the number of mitochondria in BBB endothelial cells is up to twice as high as in endothelial cells in other parts of body [60]. Likewise, the known increase in permeability of BBB as a consequence of mitochondrial dysfunction [61] is consistent with our observations of TEER reduction and permeability increase. Furthermore, other studies have established that induction of the intracellular ROS content by αSN-AGs [59] could influence BBB integrity through activation of MMPs and redistribution and degradation of TJs [62, 63]. In Fig. 9, we summarize our observations on these intracellular phenomena in the brain endothelial cells in accordance with other NVU cells.

**Fig. 9.**
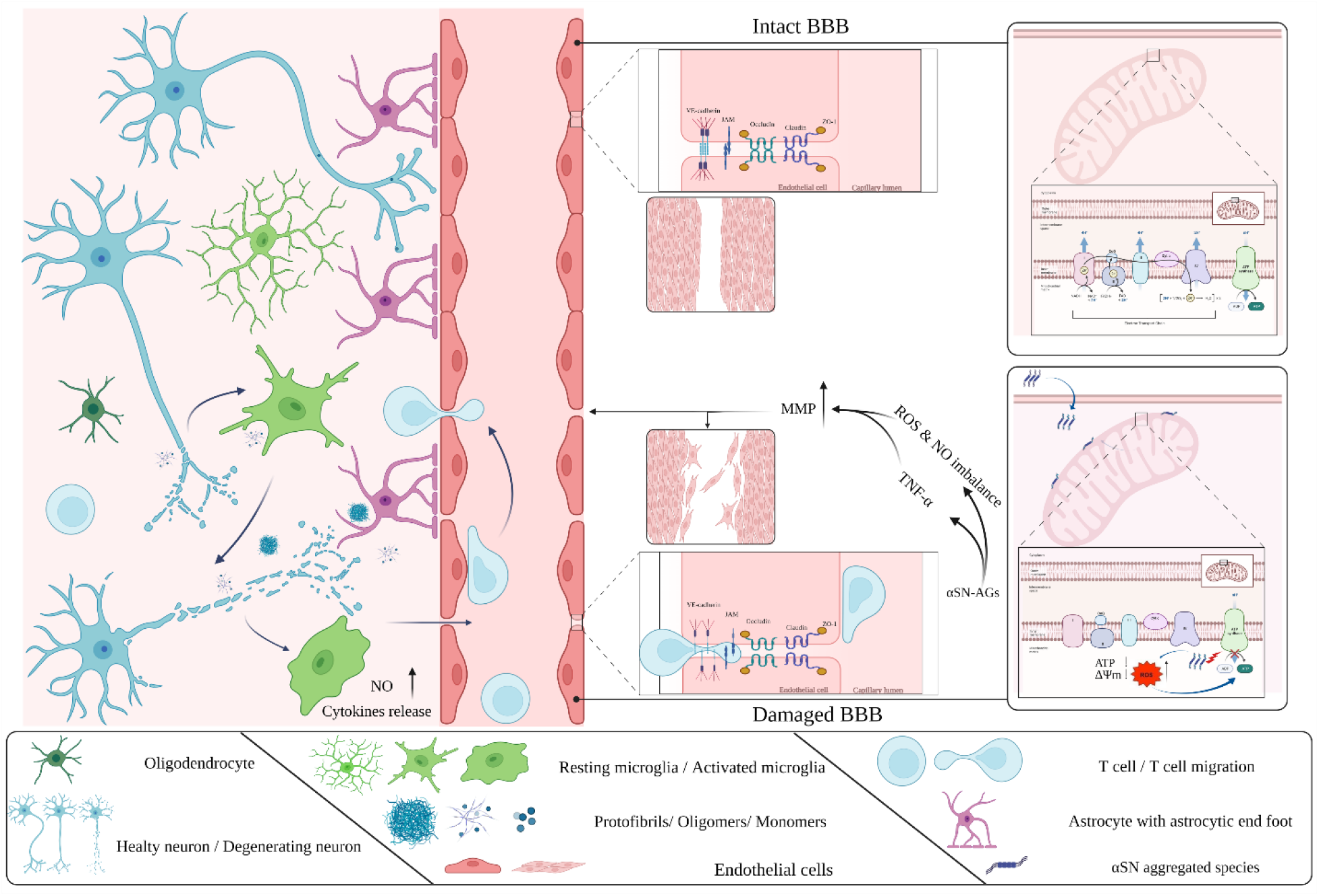
Schematic illustration of physiological and pathological characteristics of BBB during PD progression. The neurovascular unit, the basic element of the BBB, is a complex multi-cellular structure composed of endothelial cells (ECs), neurons, and glial cells (astrocytes, microglia, and oligodendrocytes) that contribute to pathological phenomena. Treatment with αSN-AGs led to a reduction of the mitochondrial membrane potential and enhancement of the intracellular ROS and NO in brain endothelial cells. Along with TNF-α expression [67], this results in wound healing impairment and BBB disruption. On the other hand, a small amount of released αSN from neurons, presumably through exocytosis, may be cleared by astrocytes or activated microglial cells. Activation of microglia may result in an inflammatory response and, as a result, neuronal degeneration, which exacerbates inflammatory conditions and immune cell recruitment, accelerating pathological progression. (Created with BioRender.com)

Regarding the impact of MMPs activation on BBB dysfunctions possibly through disruption of wound healing activity [64], it is established that they are involved in degradation of ECM-associated proteins such as proteoglycans, laminin, fibronectin, vitronectin and some types of collagens [65]. As seen in our results, treatment with αSN-AGs caused movement impairment, leading to the loss of collective cell movement in favor of single cell movement. In the collective movement pattern/sheet migration, a population of cells and their surrounding matrixes start to migrate to split sites, in order to fill and close the open area properly [66].

Increased production of pro-inflammatory cytokines (IL1β, IL6 and TNFα) in supernatant of peripheral blood mononuclear cells treated with LPS and LPS+ monomeric or oligomeric αSN, has been reported [67]. Furthermore, it has been reported that TNF-α makes a gap between endothelial cells through tyrosine phosphorylation of VE-cadherin [68], and also induces MMPs (MMP9) expression [69]. The ability of activated endothelial cells to produce pro-inflammatory cytokines such as IL1β, IL6 and TNFα [70] demonstrates a synergistic role of αSN in induction of these pro-inflammatory cytokines. Our study shows that αSN aggregates disrupt wound healing process, and we suggest that this may happen through induction of TNF-α expression in brain endothelial cells [67, 70] beyond activation of MMPs via enhancement of intracellular ROS and NO production (Fig. 9). In accordance with our results, which implicate aberrant angiogenesis in presence of αSN-AGs and endothelial disability in wound healing, cerebral microbleeds (CMBs) are more prevalent in PD patients than healthy persons [71].

### The lipid composition of cells membranes may determine how αSN interacts with the cells

For further insight into the cytotoxic mechanisms of the αSN-AGs and their impact on BBB integrity, we used FITC-αSN to show that hCMEC/D3 cells exhibit a higher propensity for αSN-AGs than HUVEC as non-BBB endothelial cells. It has been reported that αSN, especially its monomeric and oligomeric forms, shows a high interaction with anionic phospholipids through their N-terminal region [72]. The membrane binding region of αSN (N-terminal and NAC region) with hydrophobic peptide sequences, causes its penetration into the bilayer core and leads to disruption of lipid packing [73]. This disruption allows access to the bilayer membrane’s hydrophobic core and thus increases membrane permeability [72–74]. Consistent with this, our data show that brain endothelial cells, with more anionic membrane to maintain the brain homeostasis, provide lower permeability [75] but also more interaction with labeled αSN compared to HUVEC cells. Recently, the mechanism and trafficking route of αSN across BBB have been demonstrated, indicating monomeric αSN internalization via clathrin-dependent endocytosis and subsequent retromer-mediated endosome, as well as its polarized trafficking [21]. Furthermore, internalization and bidirectional transportation of αSN in monomeric and oligomeric form have been shown [21].

### BBB responses to αSN may extend beyond endothelial cells to other NVU cells with protective or attenuative roles

There is an underlying consensus that BBB is a complex system and the BBB’s endothelial cells have strong communication with other related cells including astrocytes, pericytes, neurons, and also microglial cells [11], which influence the endothelial cells in different situations, such as pathological conditions. This is schematically shown in Fig. 9. In respect to this hypothesis, we therefore further queried the effects of astrocytes, dopaminergic neurons, as well as microglial cells on the endothelial cells and their responses against αSN-AGs which highlighted the role of microglia in inflammation and deterioration of PD progression (Fig. 9).

At first, the effects of astrocytes were explored in two treatment models. In non-contact treatment model, astrocyte-conditioned media (Ast-Cond-Med) was added to hCMEC/D3 cell media while the cells were exposed to αSN-AGs. Ast-Cond-Med showed a protective effect against αSN cytotoxicity. In the second model, astrocytes were co-cultured with hCMEC/D3 in the basolateral and apical sides of inserts, respectively, and BBB function (TEER values) of the co-culture model consisting of hCMEC/D3, 1321N1 and WT SH-SY5Y, WT SH-SY5Y treated with αSN-AGs, and O-SN SH-SY5Y was affected by astrocytic protective factors (less reduction of TEER values in two groups treated with αSN).

One of the astrocyte-derived protective factors which is involved in endothelial cell survival and vascular remodeling is Angiopoietin-1 (ANG-1) that interacts with Tie-2, a tyrosine kinase receptor expressed in endothelial cells [76]. Furthermore, ANG-1 upregulated ZO-1 and occludin after ischemic damage [77]. Another astrocyte-derived factor is glial-derived neurotrophic factor (GDNF) that promotes upregulation of claudin 5 and enhancement of TEER value [78].

Additionally, by using FITC-αSN-AGs, we showed that astrocytes had a high potential to interact with αSN-AGs which could help hCMEC/D3 cells against αSN-AGs hazardous effects. Another study showed astrocytes could uptake and degrade αSN-AGs [22]. Moreover, neurons as well as astrocytes produce insulin-like growth factor-1 (IGF-1) with an anti-apoptotic role with protective effects against cell death [76, 79].

In contrast, BBB function has been affected more under inflammatory conditions, which is demonstrated by the co-culture model of BBB consisting of hCMEC/D3, 1321N1, and mixed glial cells. Different co-culturing models indicated that activated glial cells intensified αSN effects against hCMEC/D3 characteristics. Activation of microglia in response to αSN has been confirmed via NO production. NO as an inflammatory mediator, released by activated microglia (termed M1) in response to stimuli via induction of iNOS expression [80]. In the same vein and in accordance with our results, activation of microglia through TLR2 ligand activity as a result of oligomeric αSN which is released from neurons (differentiated SH-SY5Y cells overexpressing human αSN), has been revealed [24]. As seen in our results, LPS as a positive control, conditioned media collected from O-αSN SH-SY5Y cells and exogenous 12-h αSN-AGs could activate primary microglia-rich mixed glial cells. In all three groups, enhancement of NO, as well as morphological change of microglia from ramified (resting microglia) to bushy and amoeboid morphology (activated microglia), were determined.

Furthermore, the effects of glial cells have been confirmed by transmigration assay. In order to clarify the possible mechanism in the transmigration process, different studies were reviewed. It has been demonstrated that extra-cellular αSN and LPS induce IFN-γ secretion from activated microglia [81]. As well, IFN-γ has been shown to intensify αSN neurotoxicity through induction of neuro-inflammation and glial activation [82]. Activation of microglia occurs via the MyD88-dependent TLR1/2 pathway in response to oligomeric αSN [83]. The source of αSN-AGs which activate microglia, could be from apoptotic dopaminergic neurons [84] or exosomes and extracellular vehicles (EVs) or with unknown origin [85, 86]. The inflammation process begins with the activation of microglia in response to αSN-AGs, which results in the production of pro-inflammatory cytokines such as TNF-α [87, 88]. The attenuating effects of TNF-α on the impairment of BBB have been demonstrated in several studies [89–91]. Furthermore, TNF-α can participate on recruitment of T cells, by enhancing integrin ligands (such as ICAM-1, VCAM-1) on endothelial cells [92], or by increasing the propensity of T cells for adhering on the brain’s endothelial cells [93].

## Conclusion

Our data provides conclusive evidence that αSN aggregates cause endothelial cell dysregulation. Astrocytes play a protective role in co-culture models through their ability to interact with toxic αSN-AGs. Conversely, the astrocytic protective role in inflammatory conditions seems to be reversed and, in association with other glial cells, especially microglia, contributes to BBB impairment (Fig. 9). The strong and direct roles of αSN-AGs, and their role in abnormalities in vascular wound healing, leading to cerebral microbleeding, have therapeutic implications.

## Acknowledgementss

This work was supported by the Department of Industrial and Environmental Biotechnology of NIGEB. D.E.O. is grateful to the Lundbeck Foundation (grant R276-2018-671) for continued support for work on Parkinson’s Disease and to the Novo Nordisk Foundation (grant NNF17OC0028806) for previous support for work on the BBB. Graphical abstract was created with BioRender.com.

## Supplementary information

**Supplementary Figure 1.**
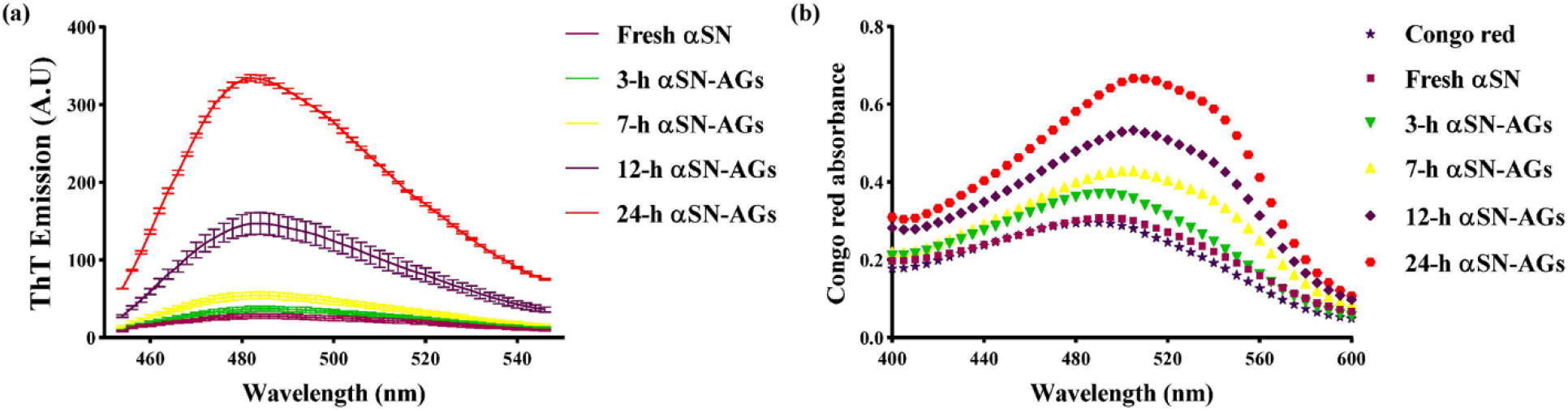
Validation of αSN-AGs. Formation of αSN-AGs after different periods of incubation using (a) ThT assay and (b) Congo red absorbance assay.

**Supplementary Figure 2.**
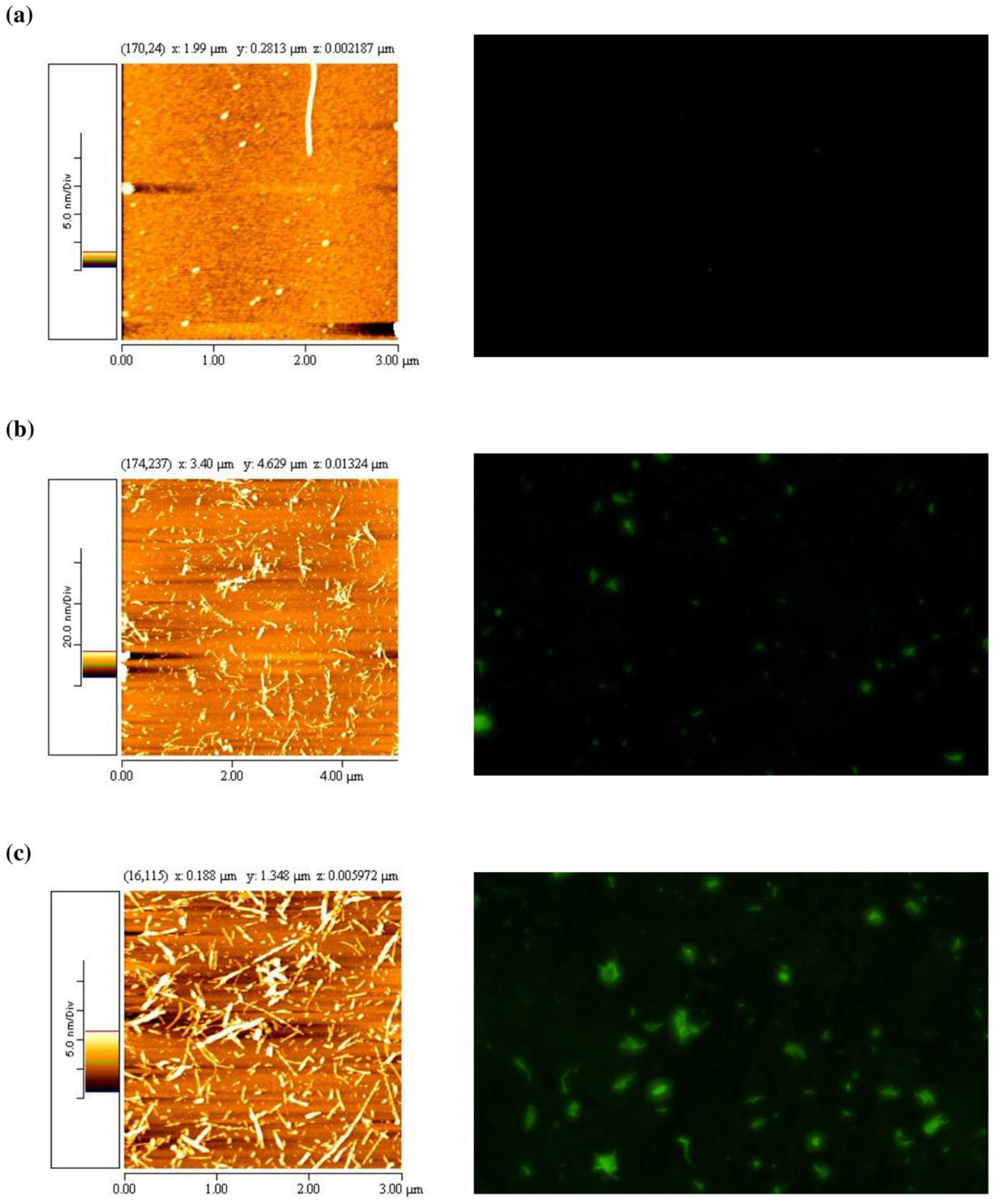
Imaging of α-SN AGs by AFM and ThT fluorescence microscopy. AFM (left) and fluorescence microscopy (right) images of 3-h α-SN AGs (a), 12-h α-SN AGs (b) and 24-h α-SN AGs (c).

**Supplementary Figure 3.**
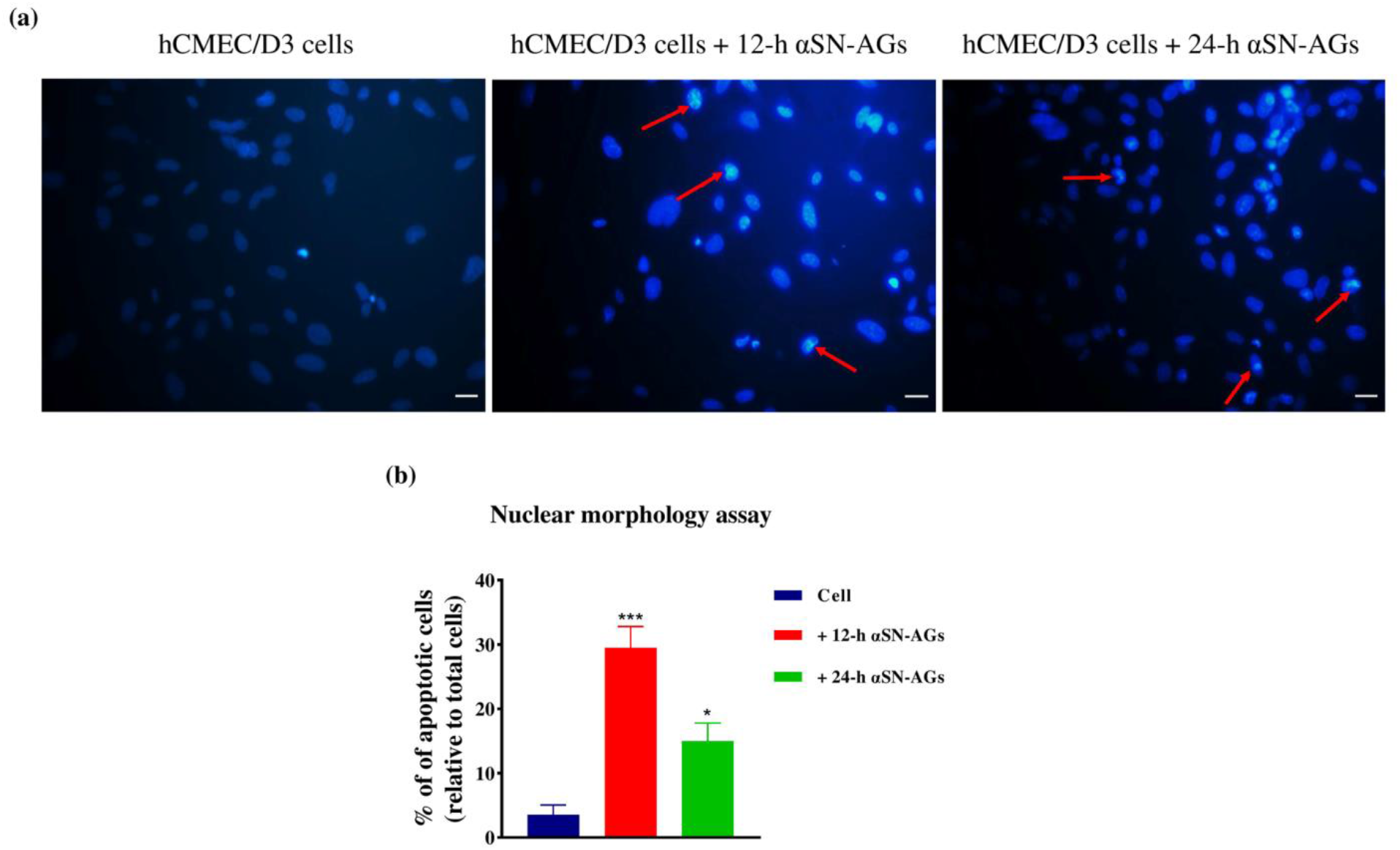
Nuclear morphology assay by DAPI representing apoptotic hCMEC/D3 cells. (a-1) hCMEC/D3 cells as control, hCMEC/D3 cells treated with (a-2) 12-h αSN-AGs and (a-3) 24-h αSN-AGs. Nuclear morphology showed apparent changes such as chromatin condensation and nuclear fragmentation (some of them displayed by red arrows) which represented apoptosis in 12-h αSN-AGs and 24-h αSN-AGs. (b) The amount of cells (%) with chromatin condensation and nuclear fragmentation. Scale bar: 20 μm.

**Supplementary Figure 4.**
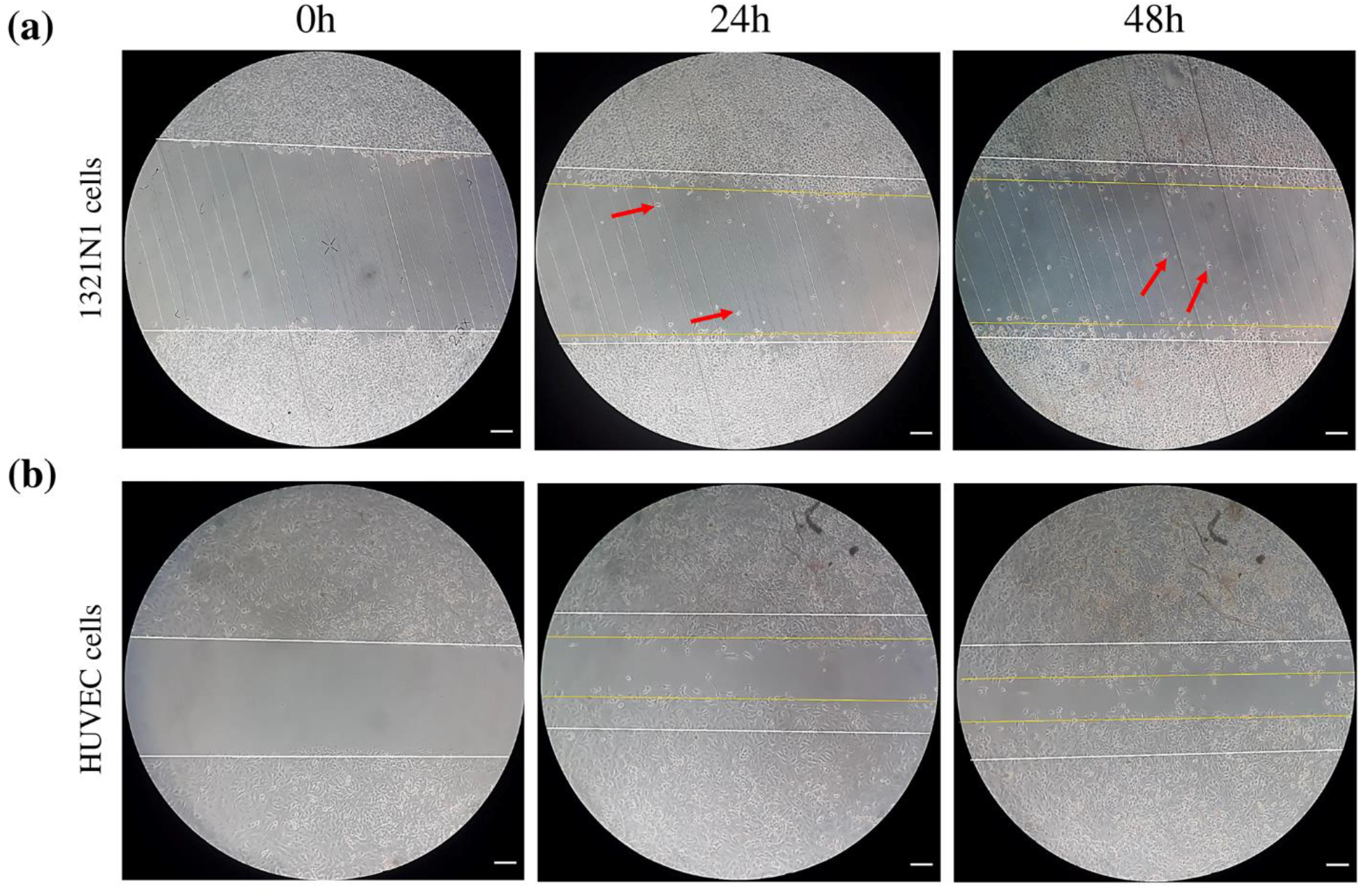
Scratch assay of HUVEC (a) and 1321N1 (b) cells visualized by phase contrast microscopy. The scratched field were assessing after 24 and 48 h using ImageJ. Single cells were shown by red arrows Scale bar: 50 μm.

**Supplementary Figure 5.**
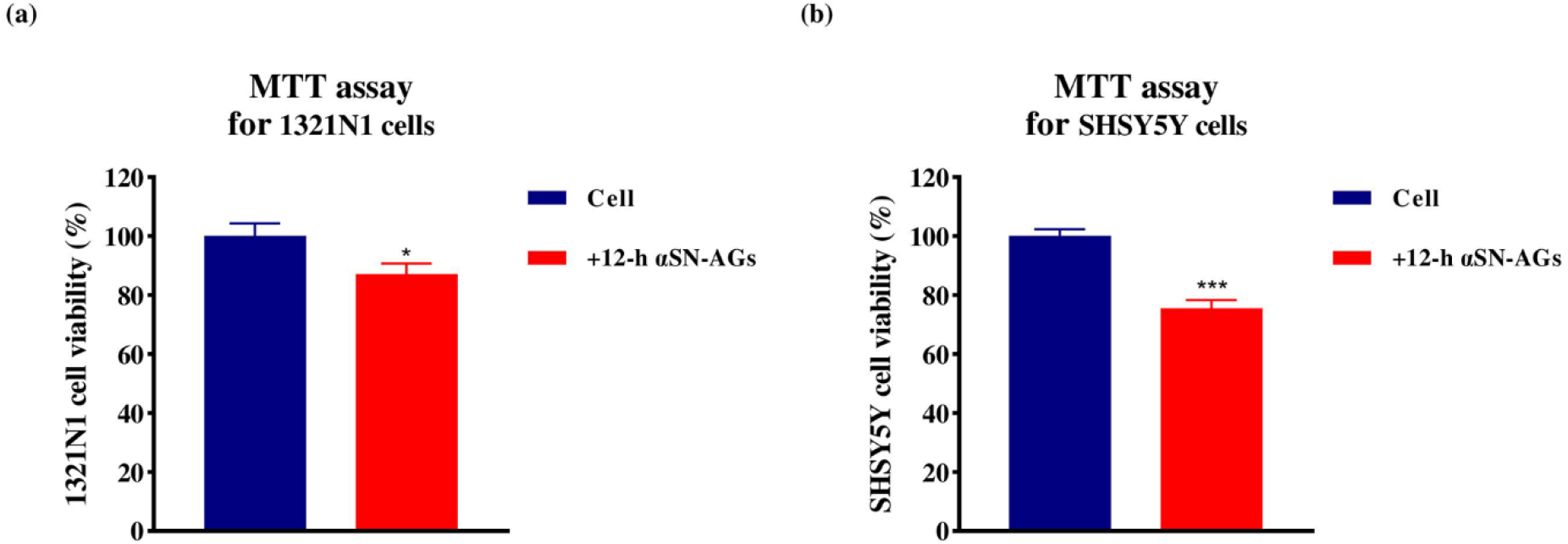
Living cell metabolic assay for 1321N1 and SHSY5Y cells treated with 12-h αSN-AGs. The cell viability of 1321N1 cells (a) and (b) SH-SY5Y cells, using MTT assay. * indicate significant differences between treated samples with that of control. (Mean±SD, * P value ≤ 0.05, *** P value ≤ 0.001).

**Supplementary Table 1:**
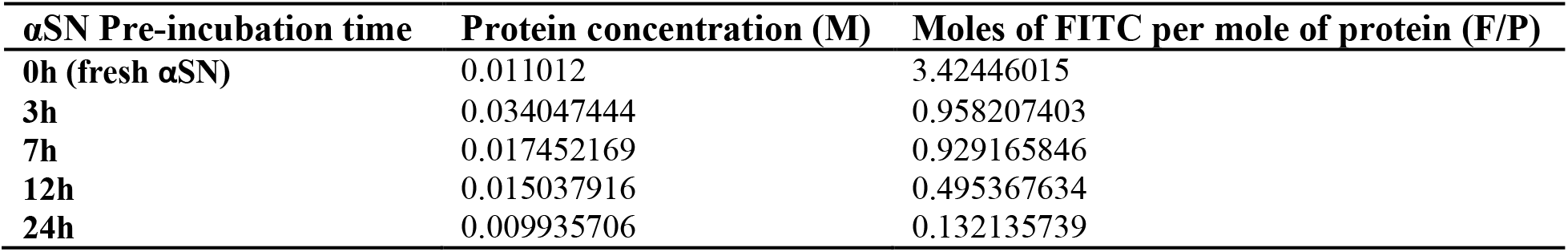
Quantification of labeled αSN. Protein concentration and moles of FITC per mole of protein (F/P) of fresh αSN and each αSN-AGs were calculated according to equations (6) and (7) in text.

| **αSN Pre-incubation time** | **Protein concentration (M)** | **Moles of FITC per mole of protein (F/P)** |
| --- | --- | --- |
| **0h (fresh αSN)** | 0.011012 | 3.42446015 |
| **3h** | 0.034047444 | 0.958207403 |
| **7h** | 0.017452169 | 0.929165846 |
| **12h** | 0.015037916 | 0.495367634 |
| **24h** | 0.009935706 | 0.132135739 |

## References

1. Kadry, H., B. Noorani, and L. Cucullo, A blood–brain barrier overview on structure, function, impairment, and biomarkers of integrity. Fluids and Barriers of the CNS, 2020. 17(1): p. 1–24.

2. Olmedo-Díaz, S., et al., An altered blood–brain barrier contributes to brain iron accumulation and neuroinflammation in the 6-OHDA rat model of Parkinson’s disease. Neuroscience, 2017. 362: p. 141–151.

3. Alvarez, J.I., et al., Focal disturbances in the blood–brain barrier are associated with formation of neuroinflammatory lesions. Neurobiology of disease, 2015. 74: p. 14–24.

4. Miners, J.S., I. Schulz, and S. Love, Differing associations between Aβ accumulation, hypoperfusion, blood–brain barrier dysfunction and loss of PDGFRB pericyte marker in the precuneus and parietal white matter in Alzheimer’s disease. Journal of Cerebral Blood Flow & Metabolism, 2018. 38(1): p. 103–115.

5. Erdő, F., L. Denes, and E. de Lange, Age-associated physiological and pathological changes at the blood–brain barrier: a review. Journal of Cerebral Blood Flow & Metabolism, 2017. 37(1): p. 4–24.

6. Gray, M.T. and J.M. Woulfe, Striatal blood–brain barrier permeability in Parkinson’s disease. Journal of Cerebral Blood Flow & Metabolism, 2015. 35(5): p. 747–750.

7. Bartels, A., et al., Blood–brain barrier P-glycoprotein function is not impaired in early Parkinson’s disease. Parkinsonism & related disorders, 2008. 14(6): p. 505–508.

8. Goedert, M., et al., 100 years of Lewy pathology. Nature Reviews Neurology, 2013. 9(1): p. 13.

9. Mahul-Mellier, A., et al., Fibril growth and seeding capacity play key roles in α-synuclein-mediated apoptotic cell death. Cell Death & Differentiation, 2015. 22(12): p. 2107–2122.

10. Alam, P., et al., α-synuclein oligomers and fibrils: a spectrum of species, a spectrum of toxicities. Journal of neurochemistry, 2019. 150(5): p. 522–534.

11. Bell, A.H., et al., The neurovascular unit: Effects of brain insults during the perinatal period. Frontiers in neuroscience, 2020. 13: p. 1452.

12. Zhang, G., et al., New perspectives on roles of alpha-synuclein in Parkinson’s disease. Frontiers in Aging Neuroscience, 2018. 10: p. 370.

13. Rocha, E.M., B. De Miranda, and L.H. Sanders, Alpha-synuclein: pathology, mitochondrial dysfunction and neuroinflammation in Parkinson’s disease. Neurobiology of disease, 2018. 109: p. 249–257.

14. Fuzzati-Armentero, M.T., S. Cerri, and F. Blandini, Peripheral-central Neuroimmune crosstalk in Parkinson’s disease: what do patients and animal models tell us? Frontiers in neurology, 2019. 10: p. 232.

15. Varatharaj, A. and I. Galea, The blood-brain barrier in systemic inflammation. Brain, behavior, and immunity, 2017. 60: p. 1–12.

16. Baltic, S., et al., α-Synuclein is expressed in different tissues during human fetal development. Journal of Molecular Neuroscience, 2004. 22(3): p. 199–203.

17. Shin, E.C., et al., Expression patterns of α-synuclein in human hematopoietic cells and in Drosophila at different developmental stages. Molecules and cells, 2000. 10(1): p. 65–70.

18. Pei, Y. and R.W. Maitta, Alpha synuclein in hematopoiesis and immunity. Heliyon, 2019. 5(10): p. e02590.

19. Shi, M., et al., Plasma exosomal α-synuclein is likely CNS-derived and increased in Parkinson’s disease. Acta neuropathologica, 2014. 128(5): p. 639–650.

20. Sui, Y.-T., et al., Alpha synuclein is transported into and out of the brain by the blood–brain barrier. Peptides, 2014. 62: p. 197–202.

21. Alam, P., et al., Polarized α-synuclein trafficking and transcytosis across brain endothelial cells via Rab7-decorated carriers. Fluids and Barriers of the CNS, 2022. 19(1): p. 1–12.

22. Booth, H.D., W.D. Hirst, and R. Wade-Martins, The role of astrocyte dysfunction in Parkinson’s disease pathogenesis. Trends in neurosciences, 2017. 40(6): p. 358–370.

23. Rannikko, E.H., S.S. Weber, and P.J. Kahle, Exogenous α-synuclein induces toll-like receptor 4 dependent inflammatory responses in astrocytes. BMC neuroscience, 2015. 16(1): p. 1–11.

24. Kim, C., et al., Neuron-released oligomeric α-synuclein is an endogenous agonist of TLR2 for paracrine activation of microglia. Nature communications, 2013. 4(1): p. 1–12.

25. Cockerill, I., et al., Blood-brain barrier integrity and clearance of amyloid-β from the BBB. Molecular, cellular, and tissue engineering of the vascular system, 2018: p. 261–278.

26. Vekrellis, K., et al., Inducible over-expression of wild type α-synuclein in human neuronal cells leads to caspase-dependent non-apoptotic death. Journal of neurochemistry, 2009. 109(5): p. 1348–1362.

27. Heravi, M., et al., The primary neuronal cells are more resistant than PC12 cells to α-synuclein toxic aggregates. Neuroscience letters, 2019. 701: p. 38–47.

28. Aliakbari, F., et al., Multiple Protective Roles of Nanoliposome-Incorporated Baicalein against Alpha-Synuclein Aggregates. Advanced Functional Materials, 2021. 31(7): p. 2007765.

29. Cabaleiro-Lago, C., O. Szczepankiewicz, and S. Linse, The effect of nanoparticles on amyloid aggregation depends on the protein stability and intrinsic aggregation rate. Langmuir, 2012. 28(3): p. 1852–1857.

30. Yakupova, E.I., et al., Congo Red and amyloids: history and relationship. Bioscience reports, 2019. 39(1).

31. Adamcik, J. and R. Mezzenga, Study of amyloid fibrils via atomic force microscopy. Current Opinion in Colloid & Interface Science, 2012. 17(6): p. 369–376.

32. Ban, T., et al., Direct observation of amyloid fibril growth monitored by thioflavin T fluorescence* 210. Journal of Biological Chemistry, 2003. 278(19): p. 16462–16465.

33. Kuan, W.-L., et al., α-Synuclein pre-formed fibrils impair tight junction protein expression without affecting cerebral endothelial cell function. Experimental neurology, 2016. 285: p. 72–81.

34. Gomes, M.J., et al., A new approach for a blood-brain barrier model based on phospholipid vesicles: Membrane development and siRNA-loaded nanoparticles permeability. Journal of Membrane Science, 2016. 503: p. 8–15.

35. Weksler, B., I.A. Romero, and P.-O. Couraud, The hCMEC/D3 cell line as a model of the human blood brain barrier. Fluids and Barriers of the CNS, 2013. 10(1): p. 1–10.

36. Parsafar, S., et al., Insights into the inhibitory mechanism of skullcapflavone II against α-synuclein aggregation and its mediated cytotoxicity. International Journal of Biological Macromolecules, 2022.

37. Perry, S.W., et al., Mitochondrial membrane potential probes and the proton gradient: a practical usage guide. Biotechniques, 2011. 50(2): p. 98–115.

38. Li, X., et al., Organic fluorescent probes for detecting mitochondrial membrane potential. Coordination Chemistry Reviews, 2020. 420: p. 213419.

39. Cummings, B.S., L.P. Wills, and R.G. Schnellmann, Measurement of cell death in mammalian cells. Current protocols in pharmacology, 2012. 56(1): p. 12.8. 1–12.8. 24.

40. Grada, A., et al., Research techniques made simple: analysis of collective cell migration using the wound healing assay. Journal of Investigative Dermatology, 2017. 137(2): p. e11–e16.

41. Bobadilla, A.V.P., et al., In vitro cell migration quantification method for scratch assays. Journal of the Royal Society Interface, 2019. 16(151): p. 20180709.

42. Srinivasan, B., et al., TEER measurement techniques for in vitro barrier model systems. SLAS Technology, 2015. 20(2): p. 107–126.

43. Heymans, M., et al., Mimicking brain tissue binding in an in vitro model of the blood-brain barrier illustrates differences between in vitro and in vivo methods for assessing the rate of brain penetration. European Journal of Pharmaceutics and Biopharmaceutics, 2018. 127: p. 453–461.

44. Guevara, I., et al., Determination of nitrite/nitrate in human biological material by the simple Griess reaction. Clinica chimica acta, 1998. 274(2): p. 177–188.

45. Paradis, A., D. Leblanc, and N. Dumais, Optimization of an in vitro human blood–brain barrier model: Application to blood monocyte transmigration assays. MethodsX, 2016. 3: p. 25–34.

46. Khalife, M., et al., Alpha-synuclein fibrils interact with dopamine reducing its cytotoxicity on PC12 cells. The protein journal, 2015. 34(4): p. 291–303.

47. Barbero, N., C. Barolo, and G. Viscardi, Bovine serum albumin bioconjugation with FITC. 2016.

48. Chaganti, L.K., N. Venkatakrishnan, and K. Bose, An efficient method for FITC labelling of proteins using tandem affinity purification. Bioscience reports, 2018. 38(6): p. BSR20181764.

49. Morshedi, D., et al., Cuminaldehyde as the major component of Cuminum cyminum, a natural aldehyde with inhibitory effect on alpha-synuclein fibrillation and cytotoxicity. Journal of Food Science, 2015. 80(10): p. H2336–H2345.

50. Crews, F.T. and R.P. Vetreno, Mechanisms of neuroimmune gene induction in alcoholism. Psychopharmacology, 2016. 233(9): p. 1543–1557.

51. De Simone, U., et al., Human co-culture model of neurons and astrocytes to test acute cytotoxicity of neurotoxic compounds. International journal of toxicology, 2017. 36(6): p. 463–477.

52. Jeon, M.-T., et al., Emerging pathogenic role of peripheral blood factors following BBB disruption in neurodegenerative disease. Ageing Research Reviews, 2021. 68: p. 101333.

53. Liang, B. and J. Su, Inducible nitric oxide synthase (iNOS) mediates vascular endothelial cell apoptosis in grass carp reovirus (GCRV)-induced hemorrhage. International journal of molecular sciences, 2019. 20(24): p. 6335.

54. Förstermann, U. and W.C. Sessa, Nitric oxide synthases: regulation and function. European heart journal, 2012. 33(7): p. 829–837.

55. Bernardini, C., et al., Differential expression of nitric oxide synthases in porcine aortic endothelial cells during LPS-induced apoptosis. Journal of Inflammation, 2012. 9(1): p. 1–9.

56. Lehner, C., et al., Oxidative stress and blood–brain barrier dysfunction under particular consideration of matrix metalloproteinases. Antioxidants & redox signaling, 2011. 15(5): p. 1305–1323.

57. Chen, X.-m., et al., Targeting reactive nitrogen species: a promising therapeutic strategy for cerebral ischemia-reperfusion injury. Acta Pharmacologica Sinica, 2013. 34(1): p. 67–77.

58. Pediaditakis, I., et al., Modeling alpha-synuclein pathology in a human brain-chip to assess blood-brain barrier disruption. Nature Communications, 2021. 12(1): p. 1–17.

59. Zadali, R., et al., A study on the interaction of the amyloid fibrils of α-synuclein and hen egg white lysozyme with biological membranes. Biochimica et Biophysica Acta (BBA)-Biomembranes, 2021: p. 183776.

60. Oldendorf, W.H., M.E. Cornford, and W.J. Brown, The large apparent work capability of the blood-brain barrier: a study of the mitochondrial content of capillary endothelial cells in brain and other tissues of the rat. Annals of Neurology: Official Journal of the American Neurological Association and the Child Neurology Society, 1977. 1(5): p. 409–417.

61. Doll, D.N., et al., Mitochondrial crisis in cerebrovascular endothelial cells opens the blood–brain barrier. Stroke, 2015. 46(6): p. 1681–1689.

62. Gu, Y., C.M. Dee, and J. Shen, Interaction of free radicals, matrix metalloproteinases and caveolin-1 impacts blood-brain barrier permeability. Front Biosci (Schol Ed), 2011. 3(3): p. 1216–1231.

63. Gu, Y., et al., Caveolin-1 regulates nitric oxide-mediated matrix metalloproteinases activity and blood–brain barrier permeability in focal cerebral ischemia and reperfusion injury. Journal of neurochemistry, 2012. 120(1): p. 147–156.

64. Sabino, F. and U. Keller, Matrix metalloproteinases in impaired wound healing. Metalloproteinases Med, 2015. 2: p. 1–8.

65. Mouw, J.K., G. Ou, and V.M. Weaver, Extracellular matrix assembly: a multiscale deconstruction. Nature reviews Molecular cell biology, 2014. 15(12): p. 771–785.

66. Vitorino, P. and T. Meyer, Modular control of endothelial sheet migration. Genes & development, 2008. 22(23): p. 3268–3281.

67. Piancone, F., et al., Inflammatory responses to monomeric and aggregated α-synuclein in peripheral blood of parkinson disease patients. Frontiers in Neuroscience, 2021. 15.

68. Angelini, D.J., et al., TNF-α increases tyrosine phosphorylation of vascular endothelial cadherin and opens the paracellular pathway through fyn activation in human lung endothelia. American Journal of Physiology-Lung Cellular and Molecular Physiology, 2006. 291(6): p. L1232–L1245.

69. Nee, L.E., et al., TNF-α and IL-1β–mediated regulation of MMP-9 and TIMP-1 in renal proximal tubular cells. Kidney international, 2004. 66(4): p. 1376–1386.

70. Pajares, M., et al., Inflammation in Parkinson’s disease: mechanisms and therapeutic implications. Cells, 2020. 9(7): p. 1687.

71. Kim, K.J., et al., The prevalence of cerebral microbleeds in non-demented Parkinson’s disease patients. Journal of Korean medical science, 2018. 33(46).

72. Iyer, A. and M.M. Claessens, Disruptive membrane interactions of alpha-synuclein aggregates. Biochimica et Biophysica Acta (BBA)-Proteins and Proteomics, 2019. 1867(5): p. 468–482.

73. Stöckl, M.T., N. Zijlstra, and V. Subramaniam, α-Synuclein oligomers: an amyloid pore? Molecular neurobiology, 2013. 47(2): p. 613–621.

74. Vasili, E., A. Dominguez-Meijide, and T.F. Outeiro, Spreading of α-Synuclein and tau: a systematic comparison of the mechanisms involved. Frontiers in molecular neuroscience, 2019. 12: p. 107.

75. Ribeiro, M., et al., Translocating the blood-brain barrier using electrostatics. Frontiers in cellular neuroscience, 2012. 6: p. 44.

76. Michinaga, S. and Y. Koyama, Dual roles of astrocyte-derived factors in regulation of blood-brain barrier function after brain damage. International journal of molecular sciences, 2019. 20(3): p. 571.

77. Xia, Y.-p., et al., Recombinant human sonic hedgehog protein regulates the expression of ZO-1 and occludin by activating angiopoietin-1 in stroke damage. PloS one, 2013. 8(7): p. e68891.

78. Shimizu, F., et al., Pericyte-derived glial cell line-derived neurotrophic factor increase the expression of claudin-5 in the blood–brain barrier and the blood-nerve barrier. Neurochemical research, 2012. 37(2): p. 401–409.

79. Wrigley, S., D. Arafa, and D. Tropea, Insulin-like growth factor 1: at the crossroads of brain development and aging. Frontiers in cellular neuroscience, 2017. 11: p. 14.

80. Orihuela, R., C.A. McPherson, and G.J. Harry, Microglial M1/M2 polarization and metabolic states. British journal of pharmacology, 2016. 173(4): p. 649–665.

81. Cebrián, C., et al., MHC-I expression renders catecholaminergic neurons susceptible to T-cell-mediated degeneration. Nature communications, 2014. 5(1): p. 1–14.

82. Wang, J., et al., Interferon-γ potentiates α-synuclein-induced neurotoxicity linked to toll-like receptors 2 and 3 and tumor necrosis factor-α in murine astrocytes. Molecular neurobiology, 2019. 56(11): p. 7664–7679.

83. Daniele, S.G., et al., Activation of MyD88-dependent TLR1/2 signaling by misfolded α-synuclein, a protein linked to neurodegenerative disorders. Science signaling, 2015. 8(376): p. ra45–ra45.

84. Bernaus, A., S. Blanco, and A. Sevilla, Glia crosstalk in neuroinflammatory diseases. Frontiers in cellular neuroscience, 2020. 14: p. 209.

85. Gustafsson, G., et al., Secretion and uptake of α-synuclein via extracellular vesicles in cultured cells. Cellular and molecular neurobiology, 2018. 38(8): p. 1539–1550.

86. Danzer, K.M., et al., Exosomal cell-to-cell transmission of alpha synuclein oligomers. Molecular neurodegeneration, 2012. 7(1): p. 1–18.

87. Rietdijk, C.D., et al., Neuronal toll-like receptors and neuro-immunity in Parkinson’s disease, Alzheimer’s disease and stroke. 2016.

88. Azam, S., et al., Regulation of toll-like receptor (TLR) signaling pathway by polyphenols in the treatment of age-linked neurodegenerative diseases: focus on TLR4 signaling. Frontiers in immunology, 2019. 10: p. 1000.

89. Cheng, Y., et al., TNFα disrupts blood brain barrier integrity to maintain prolonged depressive-like behavior in mice. Brain, behavior, and immunity, 2018. 69: p. 556–567.

90. Ge, S., et al., Human ES-derived MSCs correct TNF-α-mediated alterations in a blood–brain barrier model. Fluids and Barriers of the CNS, 2019. 16(1): p. 1–16.

91. Munoz Pinto, M.F., et al., Selective blood-brain barrier permeabilisation of brain metastases by a type-1 receptor selective tumour necrosis factor mutein. Neuro-Oncology, 2021.

92. Zhang, X., E.P. Wojcikiewicz, and V.T. Moy, Dynamic adhesion of T lymphocytes to endothelial cells revealed by atomic force microscopy. Experimental Biology and Medicine, 2006. 231(8): p. 1306–1312.

93. Li, Q., et al., Reinforcement of integrin-mediated T-Lymphocyte adhesion by TNF-induced Inside-out Signaling. Scientific reports, 2016. 6(1): p. 1–10.

